# Transient RNA dicing reprograms functional transcriptome architecture during macrophage polarization

**DOI:** 10.64898/2026.08.18.745521

**Authors:** Nira Twaik, Or Yaakov, Deema Haj Yahia, Topaz Bistritzer, Rasha Abu-Rahmah, Hodaya Turgeman, Yuval Malka

**Author notes:** These authors contributed equally (Nira Twaik, Or Yaakov).

## Abstract

The coding potential of mature mRNAs is generally considered fixed once transcription and RNA processing are complete. We previously established RNA dicing as a post-transcriptional process that generates stable, uncapped, translation-competent RNA isoforms. Here, we identify RNA dicing as a transient post-transcriptional program that remodels mature transcripts during macrophage polarization. Long-read transcriptomics reveals widespread, fate-specific dicing that peaks during early cell-state transitions and preferentially occurs between protein domains, preserving downstream coding modules. Fractionated proteomics links these RNA isoforms to truncated protein products, indicating that dicing reshapes proteomic output. Using JAK1 as a mechanistic model, we show that disruption of dicing, impairs macrophage polarization towards pro-inflammatory states. Mechanistically, a diced JH1 kinase module displayed distinct substrate preferences and alters downstream signaling relative to full-length JAK1. These findings establish RNA dicing as an adaptive layer of gene regulation that reprograms transcript architecture, expands protein functional diversity, and helps shape cell-state transitions.

## Introduction

A central assumption underlying transcriptome-level interpretation is that once an mRNA has matured, its coding architecture is largely fixed, leaving translation or degradation as its principal subsequent fate. Therefore, the mature transcriptome is widely used as a proxy for the protein-coding potential of a cell. However, accumulating evidence reveals that the relationship between the transcriptome and proteome extends far beyond this conventional linear framework ^1,2^.

Individual genes can generate protein products with distinct localizations, interaction partners, and, in some cases, opposing biological activities ^3,4^. These diversities are predicted to arise from multiple regulatory mechanisms, including alternative splicing, alternative promoter usage, RNA editing, and post-translational modifications ^4–7^. However, the mechanisms underlying the rapid reconfiguration of protein-coding potential during physiological cell-state transitions remain largely unknown.

Over the past several decades, thousands of domain-deletion and domain-mapping experiments have demonstrated that discrete protein regions can mediate remarkably distinct biological activities. Prominent examples include JAK proteins ^8,9^. The expression of different combinations of protein domains can generate products with functions that differ markedly from or even oppose those of the full-length protein, owing to altered subcellular localization, binding partners, catalytic potential, and regulatory properties. However, because these studies relied largely on engineered expression systems, the physiological relevance of such modular protein architectures remains unclear, particularly in the absence of an endogenous mechanism capable of dynamically generating such modular proteins from existing transcripts.

We recently reported a process termed RNA dicing, by which mature mRNAs can undergo additional post-transcriptional cleavage at internal alternative polyadenylation (APA) sites to generate stable, autonomous, uncapped downstream RNA fragments across thousands of genes ^10–12^. Multiple omics approaches have established these RNAs as bona fide APA cleavage products, rather than products of alternative splicing or cryptic transcription initiation. These fragments persist following transcriptional arrest and are stabilized by highly structured 5′ ends that protect them from 5′-to-3′ exonucleolytic degradation by XRN exonucleases. Their translation, in turn, is promoted by m6A modifications repositioned near the newly generated 5′ end, providing a potential framework for reconciling reports that 5′-proximal m6A promotes cap-independent translation ^13,14^, whereas internal m6A can repress translation by redirecting transcripts from polysomes to P-bodies ^15^. Thus, the functional consequences of m6A may depend on its positional context within the RNA and on the metabolic state associated with RNA dicing. We have previously established the functional consequences of this mechanism for individual genes. In BCL2, APA cleavage within the long 5′ UTR generates a stable, uncapped downstream transcript that produces full-length BCL2 through an m6A-dependent mechanism involving modification adjacent to the translation start site ^11^. Recently, we showed that dicing within the JAK1 coding sequence generates an autonomous JH1 kinase module whose activity opposes that of full-length JAK1. This truncated JH1 arises through defined APA cleavage and is translated via an m6A-dependent mechanism ^12^. Together, these studies established RNA dicing as a post-transcriptional mechanism that remodels mature transcripts into independently translated RNA units with distinct biological activities.

While these findings established RNA dicing as a broadly acting post-transcriptional mechanism that expands gene product diversity, the cellular conditions that engage this process, the biological rationale for its activation, and the extent to which it operates robustly across cellular contexts remain unclear. Notably, the core determinants underlying RNA dicing, including APA, m6A modification, RNA structure, and cap-independent translation, are dynamically remodeled across various biological transitions, including immune cell activation and differentiation, cell cycle progression, circadian regulation, and cancer ^16–18^. For example, during monocyte-to-macrophage differentiation, APA regulators, including CFIm25 (NUDT21), which we previously identified as a major regulator of RNA dicing ^11^, and CstF64, are dynamically regulated and contribute to the transition ^19,20^, while m6A metabolism is similarly remodeled ^17^. These observations raise the possibility that RNA dicing is not simply a constitutive feature of RNA metabolism but a regulated program deployed during changes in the cell state. Therefore, we investigated whether coordinated changes in RNA metabolism converge on RNA dicing to drive broader post-transcriptional and translational reprogramming.

Macrophage polarization provides a tractable model to test this hypothesis, as cells rapidly transition toward distinct pro-inflammatory (M1-like) or anti-inflammatory (M2-like) states. The mechanisms that shape the earliest phases of macrophage polarization and cell-state transition remain poorly understood ^21–23^. Moreover, because JAK1 participates in signaling pathways that preferentially drive M1-like polarization ^24–26^, it provides a tractable framework to test whether RNA dicing contributes dynamically to divergent cell-state transitions.

Here, we show that RNA dicing functions as a dynamic, transient, and actively regulated program deployed across thousands of genes during the early phase of macrophage cell-state transition. Using long-read transcriptomics, m6A mapping, size-resolved proteomics, and a dicing-deficient JAK1 allele, we demonstrate that this program is regulated rather than incidental: cleavage preferentially occurs within inter-domain coding regions, differs between M1- and M2-like polarization, and generates protein modules with activities distinct from those of their canonical proteins. Selective disruption of JAK1 dicing while preserving the canonical protein sequence impairs macrophage differentiation, establishing a functional requirement for this post-transcriptional remodeling. Finally, we show that the dynamic expression of JAK1 isoforms remodels downstream kinase signaling, giving rise to distinct signaling states with broad pathway potential. Together, these findings reveal that the mature transcriptome is not a fixed endpoint of gene expression but a modular substrate that can be dynamically reprogrammed to expand the functional output of individual genes during rapid cell-state transitions.

## Results

### RNA dicing is a transient, stage-specific program during macrophage polarization

Canonical transcriptomic and proteomic pipelines are largely designed to quantify the abundance of known genes. In standard short-read RNA-seq workflows, fragmented reads are typically aggregated into gene-level measurements, obscuring transcript-level architecture, including unique 5′ and 3′ boundaries and alternative isoforms. Therefore, two genes with identical expression levels can have entirely different functional potentials if their transcripts differ in their integrity (**Figure 1A**). This framework rests on the assumption that the coding architecture of the mature transcriptome is largely fixed. Therefore, its diversity is primarily attributed to established co-transcriptional processes, such as alternative splicing and promoter usage, with a comparatively limited capacity for further diversification at the translational level. We recently showed that RNA dicing physically partitions a transcript into two functional RNA modules: an upstream-capped RNA and an uncapped downstream RNA. This distinction is largely invisible to conventional gene expression assays, in which the identities, boundaries, and relative abundance of the resulting RNA species may be lost or misinterpreted. To evaluate dicing across macrophage differentiation, we analyzed long-read RNA-seq of HL-60 cells, a human promyelocytic leukemia cell line ^27^, that when activated with PMA can be polarized by IFN + LPS or IL4 + IL13 exposure towards pro-inflammatory M1-like or anti-inflammatory M2-like fates at three time points (12, 24, and 72 h; **Figure 1B; Figure S1A**). Each long read provides critical isoform-level information by resolving the precise 5′ and 3′ boundaries and full-length architecture of individual RNA species. Therefore, we investigated whether transcript integrity changes dynamically during macrophage polarization and whether these changes reveal a layer of RNA remodeling obscured by conventional gene-level analysis. To address this question, we stratified transcripts into three categories according to coding sequence integrity: 5′ transcripts, containing the canonical start but lacking the stop codon; full-length (FL) transcripts, containing the complete CDS; and 3′ transcripts, containing the canonical stop but lacking the start codon, the latter predominantly representing downstream products of RNA dicing. This approach enabled us to assess the coding potential of the existing transcriptome at the level of individual RNA species. Isoform composition showed pronounced and reproducible dynamics, revealing extensive remodeling of the modular architecture of the transcriptome (**Figure 1C**). The 5′ class remained at a relatively steady ratio, whereas canonical FL reads ∼85% of the total in the untreated state - fell to 76% after PMA and declined further upon induction, more strongly in M1-like cells (69% vs. 68% for M2-like cells at peak). The two polarization states exhibited distinct kinetics, with M1-like dicing peaking at ∼12 h after induction and M2-like dicing at ∼24 h (69% and 68%, respectively). Strikingly, by 72 h, the response markedly declined in both polarization states, approaching PMA-only levels (76% and 75%), although residual changes persisted, particularly in M2-like cells. These kinetics indicate that dicing remodeling is predominantly transient rather than constitutive. The 3′ class mirrored this pattern, rising with PMA and again upon induction (7%→15%→20%/18% in M1-like and M2-like cells, respectively, at 12 h) before returning by 72 h. Two lines of evidence suggest that these isoforms represent regulated cleavage products rather than nonspecific degradation intermediates. First, bulk mRNAs typically persist for hours, whereas exonucleolytic decay intermediates are rapidly cleared from the cytoplasm. Because cells were collected at defined time points and immediately lysed and frozen, such intermediates are expected to remain near background steady-state levels. In contrast, we detect an abundant and readily quantifiable population of 3′ isoforms. Second, the observed fragments are unlikely to arise from post-lysis RNA degradation. Such degradation should occur stochastically and broadly across transcripts and samples, whereas the 3′-isoform landscape is gene-specific, varies systematically across conditions and time points, and produces clear, reproducible, treatment-dependent segregation by principal component analysis (**Figure S1B**). Together, these features are inconsistent with random RNA breakdown and instead support regulated transcript cleavage. In addition, we have previously shown that these downstream fragments are stable, autonomous, and exonuclease-resistant RNA isoforms ^10^. In contrast, the relative instability of the upstream 5′ class is consistent with clearance of CDS-cleaved transcripts by non-stop-mediated decay ^28^, whereas the downstream 3′ products retain the canonical in-frame stop codon. Moreover, the permissive use of non-canonical translation initiation codons, including NUG codons ^29,30^, enables these downstream fragments to retain substantial coding potential despite the loss of the canonical start codon.

**Figure 1.**
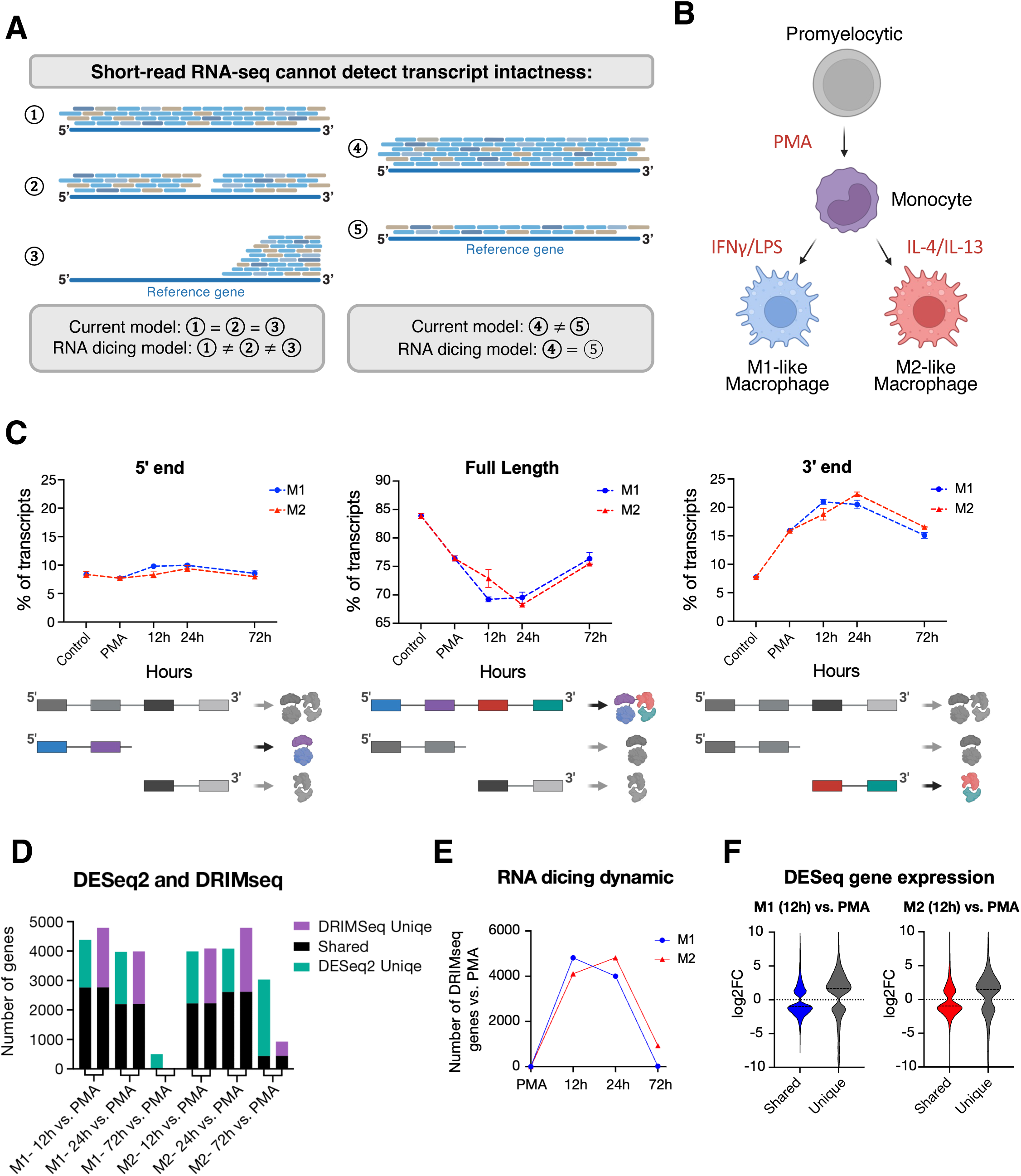
RNA dicing is a transient, stage-specific program during macrophage polarization. **(A)** Schematic illustrating how conventional short-read RNA-seq can obscure transcript intactness. Transcripts with similar gene-level read coverage can represent full-length or independently generated 5′ and 3′ RNA species (DESeq2 analysis), whereas changes in transcript composition can also alter the relative abundance of specific diced isoforms. **(B)** Experimental framework for HL-60 differentiation. Promyelocytic HL-60 cells were treated with PMA and subsequently polarized toward M1 macrophages with IFNγ/LPS or M2 macrophages with IL-4/IL-13. **(C)** Relative abundance of 5′, full-length (FL), and 3′ transcript classes across control, PMA, and M1- or M2-polarized cells at 12, 24, and 72 h. Loss of FL transcripts is accompanied by a reciprocal increase in 3′ isoforms during early polarization, followed by a return toward the PMA state. **(D)** Numbers of genes identified as significantly altered by DESeq2 and DRIMSeq across M1 and M2 time points relative to PMA, showing genes uniquely detected by each method and genes shared between the two analyses. **(E)** Temporal dynamics of differentially diced genes identified by DRIMSeq relative to PMA during M1 and M2 polarization. **(F)** Distribution of DESeq2 log2 fold changes for genes detected by both DESeq2 and DRIMSeq (“shared”) or uniquely by DESeq2 at 12 h of M1 and M2 polarization, illustrating how extensive transcript remodeling can influence conventional gene-level abundance estimates.

To distinguish normalized bulk gene expression from differential expression of non-canonical RNA modules, we analyzed the data using two complementary statistical frameworks. DESeq2, the standard for differential gene expression, quantifies total gene-level abundance across conditions and time points ^31^. Importantly, standard differential abundance analyses fail to capture shifts in the relative proportions of transcript isoforms (**Figure 1A**). To capture these dicing-driven events, we employed DRIMSeq, which models transcript usage via a Dirichlet-multinomial distribution, to test for isoform composition changes independently of total gene expression ^32^. Across both DESeq2 and DRIMSeq frameworks, intermediate time points (12 and 24 h) shared ∼60% of the significant genes (**Figure 1D**). By 72 h, the two analyses revealed markedly different trajectories. DESeq2 showed persistent separation of M1-like and M2-like cells into distinct clusters, reflecting sustained differences in overall gene abundance. In contrast, DRIMSeq showed both conditions shifting back toward the baseline PMA state, consistent with dicing-mediated isoform remodeling being predominantly transient and resolved over time (**Figure 1C**). Because the overall dicing dynamics appeared transient, we tested each time point against PMA, which served as an anchor for the start of differentiation. In M1-like cells, DRIMSeq identified 4,813 differentially diced genes at 12 h and 4,004 at 24 h, but only 13 at 72 h, revealing a pronounced yet transient wave of transcript remodeling. DESeq2 reported comparable numbers at 12–24 h; however, at 72 h, it still identified 497 differentially expressed genes (**Figure 1D-E**). A similar trend was observed in M2-differentiated cells, with a higher number of genes at 72 h, consistent with their slower differentiation rate ^33^. These results reveal two parallel and largely independent modes of transcriptome adaptation: RNA dicing, a fast catabolic response to the differentiation signal whose effect on M1-like cells is essentially fully transient and resolves by 72 h, and anabolic canonical expression change (DESeq2), a slower program in which hundreds of genes shift their expression levels and remain altered at 72 h.

Next, we evaluated the different sets of genes detected by each method. DESeq2 and DRIMSeq shared a substantial number of significant genes, yet thousands of genes were unique (**Figure 1D**). Since dicing is a catabolic process that reduces overall read coverage per gene (RPKM), we reasoned that many genes scored as downregulated by DESeq2 may, in fact, represent transcripts undergoing extensive dicing (**Figure 1A**, panel 3). Analysis of genes shared between the DESeq2 and DRIMSeq datasets supports this interpretation: a substantial fraction of genes classified as downregulated by DESeq2 instead showed a shift toward diced isoforms, indicating transcript remodeling rather than transcriptional repression or nonspecific RNA degradation (**Figure 1F**). More broadly, this compositional shift also complicates the interpretation of genes classified as uniquely upregulated by DESeq2. Because normalization is performed relative to the overall RNA population, widespread dicing-dependent loss of reads from a large fraction of transcripts could cause genes with relatively stable absolute abundance to appear increased. Thus, during extensive transcript remodeling, conventional gene-level normalization may capture both true expression changes and relative abundance shifts generated by the dicing process itself.

### Dicing is fate-specific and directed toward inter-domain sites

While macrophage identity following polarization is well-defined at later stages, the mechanisms that shape the earliest phases of cell-state transition remain poorly characterized ^21,22^. We reasoned that post-transcriptional catabolic reprogramming by RNA dicing might provide mechanistic insight into the ignition phase of differentiation. The early and transient nature of this response suggests that macrophages enter polarization with a preexisting transcriptome that can be remodeled by RNA dicing to generate fate-specific outputs before the establishment of sustained de novo transcriptional programs. Conceptually, this implies that the canonical transcriptome contains stratified layers of regulatory information that can be selectively activated through distinct dicing programs in response to diverse environmental stimuli.

We therefore asked whether M1- and M2-like macrophages diverge in their dicing programs from the earliest stages of polarization, prior to the establishment of stable phenotypic identities. To identify the earliest divergence separating pro- and anti-inflammatory states, we performed pairwise DESeq2 and DRIMSeq comparisons between M1- and M2-like cells at each time point. At the initial polarization at 12 h, both frameworks revealed substantial lineage-specific differences, with approximately 300 unique genes identified in each analysis, which rapidly contracted by 24 h (∼50 DESeq2 and ∼100 DRIMSeq hits remaining). By 72 h, as cells approached terminal differentiation, DESeq2 detected a substantial surge in differential gene expression involving thousands of genes, alongside approximately 800 DRIMSeq-specific dicing shifts (**Figure 2A**). This divergence seemed to be driven by M2-like cells maintaining active dicing, while M1-like cells returned to baseline PMA levels. Next, we analyzed consecutive time point transitions within each lineage to assess how RNA dicing evolved over the course of polarization. Both M1- and M2-like paths displayed a biphasic pattern: an immediate, rapid burst of differential abundance and dicing from 0h (PMA) to 12 h, followed by a period of minimal transcriptomic change between 12 and 24 h. As cells progressed toward terminal differentiation at 72 h, a second major transition occurred, with thousands of genes changing significantly relative to 24 h and shifting back toward the basal state, consistent with the patterns observed in **Figures 1D-E** and **Figure S2A**. MSigDB Hallmark gene set analysis revealed lineage-specific RNA-dicing programs in M1- and M2-polarized cells, including mTORC1 signaling and RNA-splicing pathways, with divergent dicing patterns that were largely invisible to conventional DESeq2-based differential expression analysis (**Figure 2B; Figure S2B**). This divergence suggests that RNA dicing is a directed, signal-specific regulatory mechanism, rather than a generic consequence of cellular activation. We next focused on two major functional protein families, protein kinases and RNA-binding proteins, which recapitulated the global dicing dynamics. Although the overall signatures were largely shared between fates, their magnitude and timing differed: dicing and 3′-isoform expression were strongest early in M1-like cells (12 h) but peaked later in M2-like cells (24 h) (**Figure 2C; Figure S2C**). Although the dicing signature provides a robust measure of the direction of transcript remodeling, it does not resolve the coding potential of the resulting isoforms or the protein products that they may generate. Therefore, we compared the average length of 3′ diced isoforms across time points in M1- and M2-like macrophages. At 12 h, M1-like cells displayed shorter 3′ isoforms compared to M2-like cells. This trend was reversed at 24 and 72 h, when shorter diced isoforms predominated in M2-like cells. Thus, the extent of transcript truncation is dynamically regulated in both time- and fate-dependent manners. (**Figure 2D; Figure S2D**).

**Figure 2.**
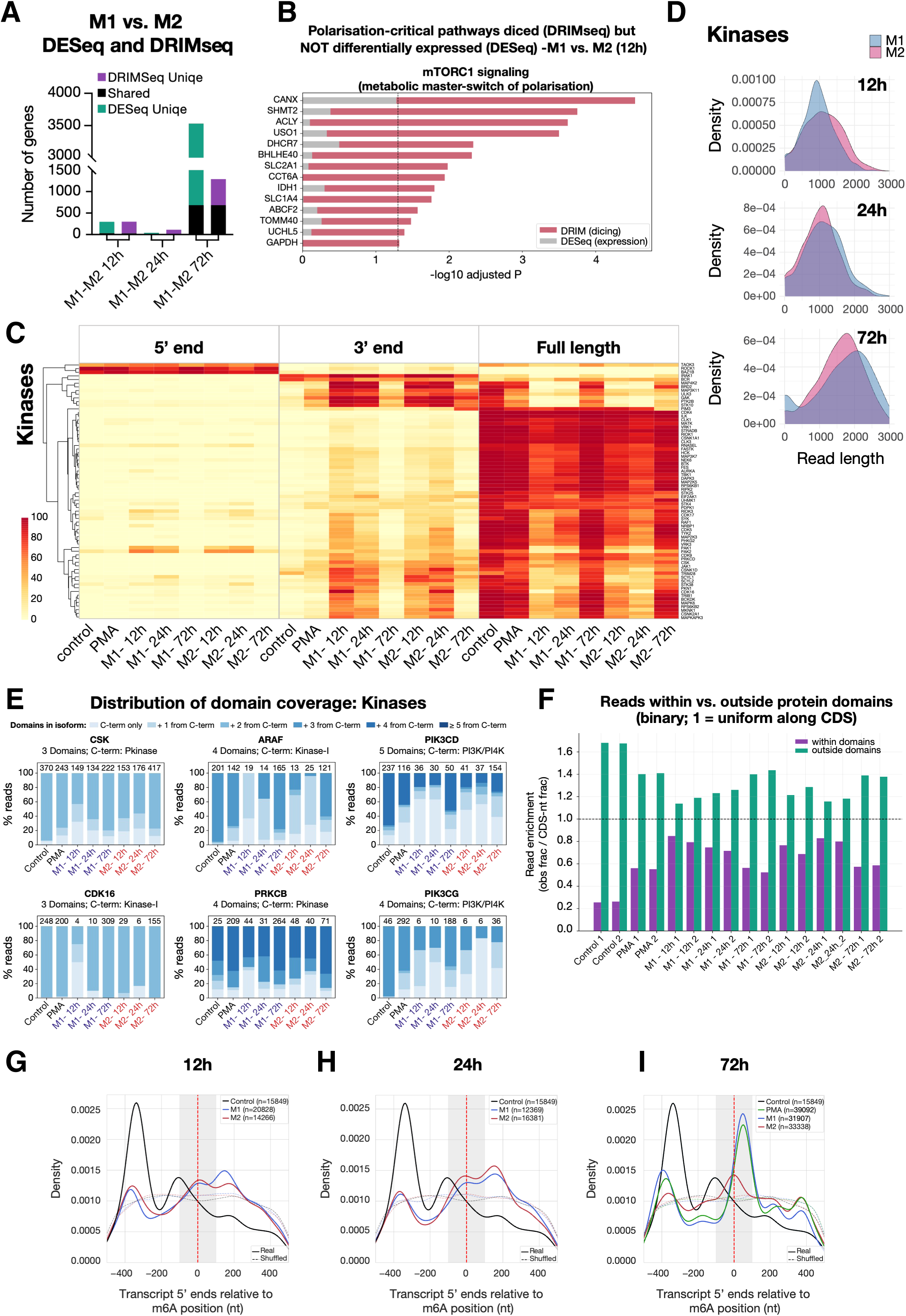
RNA dicing is fate-specific and spatially coordinated with protein-domain architecture. **(A)** Numbers of genes differing between M1 and M2 cells at matched time points (12, 24, and 72 h; upper panel) and across consecutive time-point transitions within each lineage (lower panel), as detected by DESeq2 and DRIMSeq, showing method-specific and shared events. **(B)** Differential dicing of genes associated with mTORC1 signaling at 12 h of M1 versus M2 polarization. DRIMSeq significance is compared with conventional DESeq2 differential-expression signals, highlighting pathway-associated transcript remodeling that is not captured by gene-level abundance analysis. **(C)** Heatmap of 5′, 3′, and full-length transcript classes for protein kinases across control, PMA, and M1/M2 polarization, illustrating dynamic dicing signatures and fate-dependent remodeling of kinase transcripts. **(D)** Length distributions of kinase-derived 3′ isoforms in M1 and M2 cells at 12, 24, and 72 h, revealing time- and fate-dependent differences in the extent of transcript truncation. **(E)** Enrichment of dicing-associated transcript boundaries within versus outside protein-domain coding regions, normalized to the expectation for a uniform distribution across the CDS. Dicing is preferentially enriched outside annotated domains, consistent with cleavage within inter-domain regions. **(F)** Domain coverage of representative multidomain kinases across differentiation states. Stacked bars indicate the fraction of reads retaining progressively larger numbers of domains counted from the C terminus, illustrating condition-dependent generation of transcripts with distinct domain compositions. **(G–I)** Distribution of transcript 5′ ends relative to m6A-enriched positions at 12 h (G), 24 h (H), and 72 h (I). Solid lines represent observed distributions and dotted lines shuffled controls; increased coincidence between m6A and dicing-generated neo-5′ ends.

Because long-read sequencing preserves the complete structure and translational potential of individual RNA isoforms, we next asked how dicing alters their encoded protein-domain architecture. For each 3′ isoform, we determined which annotated protein domains remained intact downstream of the neo-5′ end of the gene. This analysis revealed distinct isoform-length classes that varied with the dicing intensity (**Figure 2E; Figure S2E**). Diced transcripts typically lose upstream coding regions while retaining the complete sequence of the first downstream protein domain in a potentially translatable configuration, consistent with the generation of modular protein outputs. This pattern may suggest that RNA dicing extends beyond the generation of a single defined isoform per gene, creating a diverse repertoire of catabolic isoforms that encode distinct combinations of protein domains. Therefore, we investigated whether RNA dicing occurs randomly across the coding sequence or is spatially organized relative to the corresponding protein domain architecture. For that, we analyzed the percentage of reads that contained the number of domains upstream to the C-terminal, across conditions and time points. The cleavage sites defining the 5′ boundaries of downstream 3′ diced isoforms were preferentially located within the inter-domain coding regions (**Figure 2F; Figure S2F**). This positional bias suggests that dicing preferentially partitions transcripts between protein domains, thereby preserving intact downstream domain architecture.

We previously showed that endonucleolytic cleavage and the generation of uncapped downstream RNA molecules are insufficient for translation. Rather, this process requires an additional layer of RNA regulation, in which m6A modification of the newly generated 5′ end licenses cap-independent translation of the diced isoform ^11,13,14^. We previously proposed that dicing passively reposition internal CDS m6A modifications to the neo-5′ end of the resulting uncapped fragment, where they may acquire translation-initiation activity and promote cap-independent translation ^11,13,14^. We therefore integrated m6A-RIP data from human monocytes ^34^ with long-read RNA sequencing to determine whether m6A modifications coincide with dicing-generated neo-5′ ends. In untreated monocytes, the dominant 5′-end population was located several hundred nucleotides upstream of m6A-enriched sites, a configuration predicted to be unfavorable for m6A-mediated cap-independent translation (**Figure 2G**). at 12–24 h of M1- and M2-like cells, m6A increasingly coincided with the neo-5′ ends of diced isoforms (**Figure 2H**), reaching its highest levels in PMA and at 72 h of M1-like macrophages (**Figure 2I**). Dicing and m6A repositioning thus provide a mechanism by which thousands of truncated isoforms may gain translational competence.

Together, this positional selectivity supports a model in which RNA dicing functions as a modular, rather than degradative, process that is spatially coordinated with the domain architecture of encoded proteins.

#### RNA dicing reshapes protein isoform composition

Our previous studies established genome-wide that diced isoforms undergo m6A-dependent, cap-independent translation, as evidenced by strong polysome enrichment, mapped ribosomal initiation sites, and isoform-specific N-terminal peptides detected by mass spectrometry ^11^. These findings suggest that dicing-dependent RNA reprogramming has the potential to substantially reshape the proteomic landscape. However, conventional mass spectrometry workflows share a limitation analogous to that of short-read RNA sequencing, which provides limited information about the precise N- and C-terminal boundaries of non-canonical isoforms. Moreover, although the N-terminal proteomics approaches we previously employed enabled high-resolution mapping of dynamically regulated translation initiation sites in diced isoforms, they did not resolve the full-length translational products generated from these transcripts.

We therefore performed size-resolved differential proteomics: lysates were separated by SDS-PAGE into three fractions (10–40, 40–80, and 80–180 kDa) and analyzed by MS (**Figure 3A**). The canonical mechanisms underlying isoform diversity are alternative splicing and alternative promoter usage. In many cases, alternative splicing modifies individual exons or short transcript segments, whereas alternative promoter usage changes transcription start site selection and often alters the 5′ untranslated region or N-terminus of the encoded protein. Nevertheless, our model suggests that RNA dicing preferentially removes entire protein domains while preserving intact downstream domain architecture. We first analyzed peptides derived from intrinsically small proteins (up to 40 kDa), which showed no fractionation bias between control HL-60 cells and the 12 h time point of M1-like cells (**Figure 3B**). Next, we investigated whether peptides derived from high-molecular-weight proteins (>80 kDa) were partitioned into their expected size fractions. In control cells, these peptides were predominantly detected in the 80–180 kDa fraction, consistent with their canonical protein size. In M1-like cells, however, the same peptide population shifted toward the light (10–40 kDa) fraction, indicating that sequences derived from high-molecular-weight proteins were incorporated into substantially smaller protein isoforms. Finally, at 72h time point, we detected partial restoration of these proteins to their canonical size. A similar trend was observed at 12 and 72 h in M2-like cells (**Figure 3B**), consistent with the translational potential of diced transcripts observed across the transcriptomes. Similar to RNA turnover, protein degradation occurs on a much faster timescale than the typical half-life of intact proteins, generating short-lived, heterogeneous peptide products ^35^. Thus, the reproducible accumulation of peptides from high-molecular-weight proteins within defined low-molecular-weight fractions is unlikely to reflect non-specific protein degradation. Next, we quantified gene-level expression per fraction, and the heavy-to-light ratio showed a graded, dicing-matched shift with a temporal lag, peaking toward 24 h in M1-like cells, consistent with protein following mRNA (**Figure S3A**), whereas M2-like cells shifted at an earlier time point (**Figure S3B**). To test whether dicing is causally linked to the production of truncated isoforms, we related the heavy-to-light shift of each gene to its corresponding dicing signature. We therefore selected two groups of genes for comparison: genes uniquely significant by DESeq2, representing non-diced transcripts, and genes significant by both DESeq2 and DRIMSeq, representing diced transcripts. Genes with a dicing signature showed an increased representation of truncated protein isoforms, whereas this pattern was markedly reduced among non-diced genes (**Figure 3C-D**). In addition, genes with high dicing intensity exhibited a greater shift from heavy to light protein fractions than genes with intermediate or low dicing intensity. Together, these findings establish a strong correspondence between transcript-level dicing and the emergence of truncated protein products, supporting a model in which diced mRNA isoforms are translated into distinct and truncated proteomic outputs.

**Figure 3.**
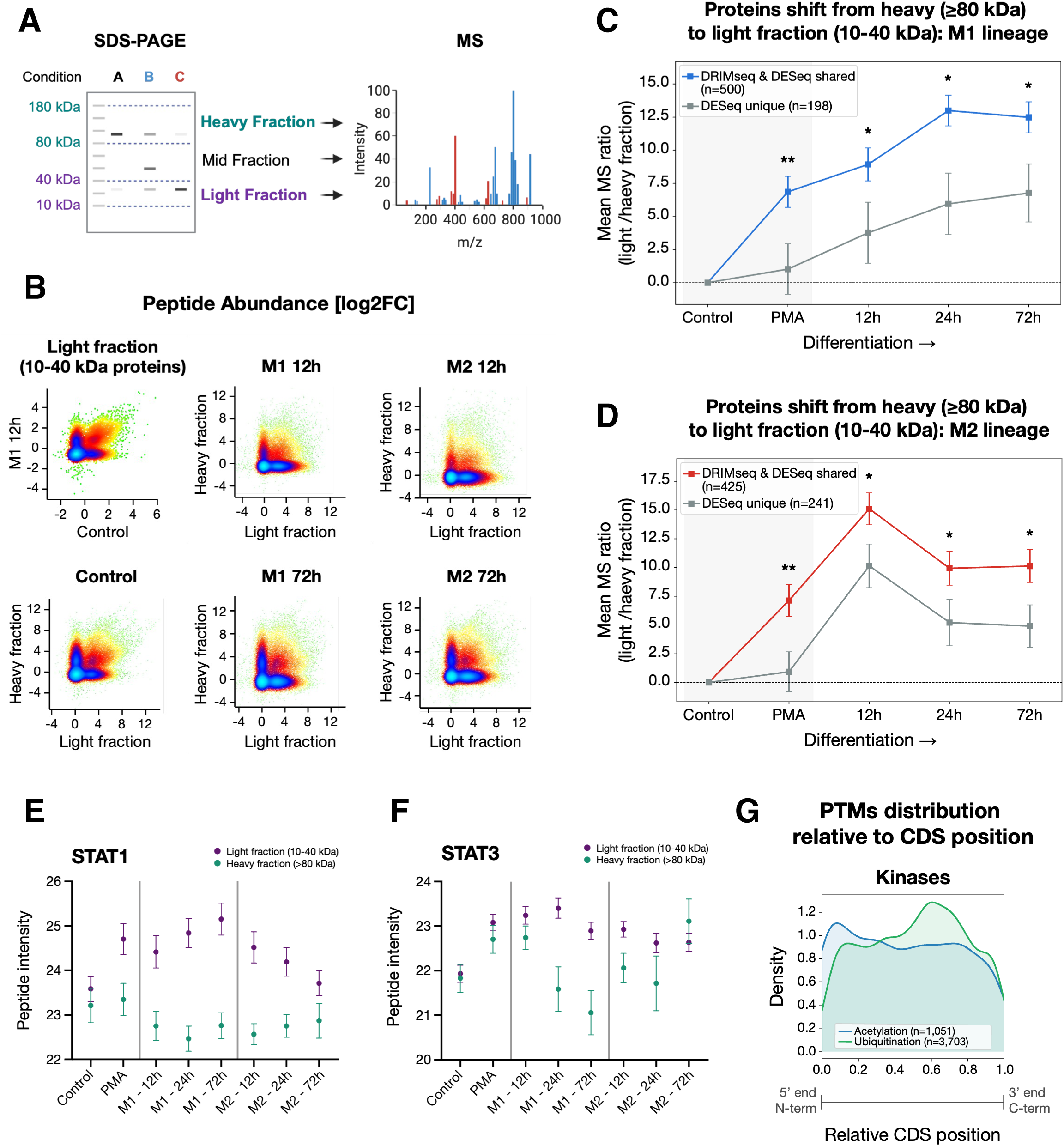
Size-resolved proteomics links RNA dicing to truncated protein isoforms. **(A)** Schematic of the size-resolved proteomics workflow. Cellular proteins were separated by SDS-PAGE into light (10–40 kDa), intermediate (40–80 kDa), and heavy (80–180 kDa) fractions, followed by mass-spectrometry analysis. **(B)** Peptide-abundance distributions across molecular-weight fractions. Naturally small proteins (< 40 kda) serve as a fractionation control, whereas peptides derived from higher-molecular-weight proteins (> 80 kDa) redistribute toward the light fraction during macrophage polarization and partially return toward their canonical size distribution at later time points. **(C, D)** Mean light-to-heavy MS ratio for genes identified by both DRIMSeq and DESeq2 compared with genes uniquely identified by DESeq2 during M1 (E) and M2 (F) polarization. Genes carrying a dicing signature show a greater shift toward low-molecular-weight protein species. **(E, F)** Size-resolved peptide intensity of STAT1 (E) and STAT3 (F) across control, PMA, and M1/M2 polarization. Light- and heavy-fraction signals reveal distinct temporal and lineage-specific changes in STAT protein isoform composition. Data are shown as mean ± SD. Statistical significance between the two gene groups at each time point was assessed using a two-tailed Student’s *t*-test; *p* < 0.05 (\**) and p < 0.01* (**) **(G)** Density distributions of acetylation (blue) and ubiquitination (green) sites along the relative coding sequence (CDS) position (0 = 5′/N-terminus, 1 = 3′/C-terminus). Each site is plotted individually; density was estimated using kernel density estimation (KDE) with Scott’s bandwidth rule.

Next, we investigated whether size-resolved proteomics could reveal the dynamic isoform regulation of individual proteins within pathways relevant to macrophage polarization. Specifically, we focused on STAT1 (91 kDa) and STAT3 (89 kDa), which are the major transcriptional effectors of JAK signaling that are preferentially associated with M1- and M2-like polarization, respectively ^36,37^. At the RNA level, both genes exhibited dynamic lineage-dependent shifts between canonical and diced isoforms during macrophage polarization. (**Figure S3D**). Therefore, we examined whether these transcript-level dynamics were reflected in the protein output. Both STAT1 and STAT3 showed redistribution between full-length (FL; 80–180 kDa) and lower-molecular-weight diced (D; 10–40 kDa) protein species, but with distinct temporal and fate-specific patterns of redistribution. For STAT1, the diced species progressively increased during M1-like polarization, whereas the abundance of the full-length protein changed modestly (**Figure 3E**). STAT3 followed a different trajectory: the full-length species progressively decreased during M1-like polarization, whereas in M2-like cells, it reached its highest abundance at 72 h (**Figure 3F**). Thus, STAT1 and STAT3 undergo distinct isoform remodeling during macrophage polarization, linking transcript-level dicing to dynamic changes in protein isoform composition over time and cell fate. Notably, alternative RNA processing of STAT family members can generate C-terminally truncated isoforms with regulatory properties that are distinct from those of their full-length counterparts. STAT1β and STAT3β, for example, retain core signaling and DNA-binding functions while lacking substantial portions of their C-terminal transactivation domains and have been reported to modulate the activity of the corresponding full-length STAT proteins ^38,39^. These findings suggest that diced STAT isoforms may reshape rather than simply suppress JAK–STAT signaling.

Finally, we asked whether the positional organization of protein domains is accompanied by a corresponding organization of post-translational modifications. For example, kinase domains are known to be preferentially positioned toward the C-termini of protein kinases (**Figure S3E**). This potentially creates an architecture in which RNA dicing could remove variable upstream regions while retaining the catalytic module. We further examined whether regulatory modifications are similarly distributed along the protein sequence. Mapping the annotated post-translational modifications in kinases from dbPTM revealed an interesting positional asymmetry: acetylation sites were preferentially enriched toward the N-terminus, whereas ubiquitination sites showed a more C-terminal distribution (**Figure 3G**). Given the established interplay between acetylation and ubiquitination in regulating protein stability ^40^, this positional organization has an additional regulatory consequence: removal of specific protein domains necessarily eliminates the post-translational modification sites and regulatory elements embedded within them, thereby altering the regulatory potential of the resulting isoforms.

### Dicing of JAK1 is required for M1-like, pro-inflammatory macrophage polarization

The results above suggest that RNA dicing regulates gene output by altering the translational potential of mature transcripts and, consequently, the repertoire of protein isoforms they produce. Features such as alternative polyadenylation site usage, m6A modification, RNA-binding protein interactions, and RNA structure collectively contribute to this regulatory architecture and influence the dicing and metabolic processing of mature mRNAs. Our previous detailed characterization of JAK1 dicing and its functional consequences in signaling pathways that preferentially drive M1-like polarization ^24–26^, makes JAK1 a particularly informative system to test this principle. The N-terminal FERM and SH2 domains of JAK1 mediate receptor interactions, followed by the regulatory JH2 pseudokinase domain and C-terminal catalytic JH1 kinase domain. We previously showed that a major JAK1 dicing event generates a truncated transcript that only encodes the JH1 domain, thereby separating the catalytic kinase from its upstream regulatory regions, including the inhibitory JH2 domain ^12^. This diced ‘JH1 module’ is translated as an autonomous ∼38-kDa protein, preferentially localizes to the nucleus, and displays enhanced kinase activity relative to full-length JAK1^12^. In vitro, the isolated JH1 module phosphorylates STAT substrates substantially more efficiently than canonical JAK1, demonstrating that dicing generates a functionally distinct kinase module rather than simply a truncated protein (**Figure 4A**).

**Figure 4.**
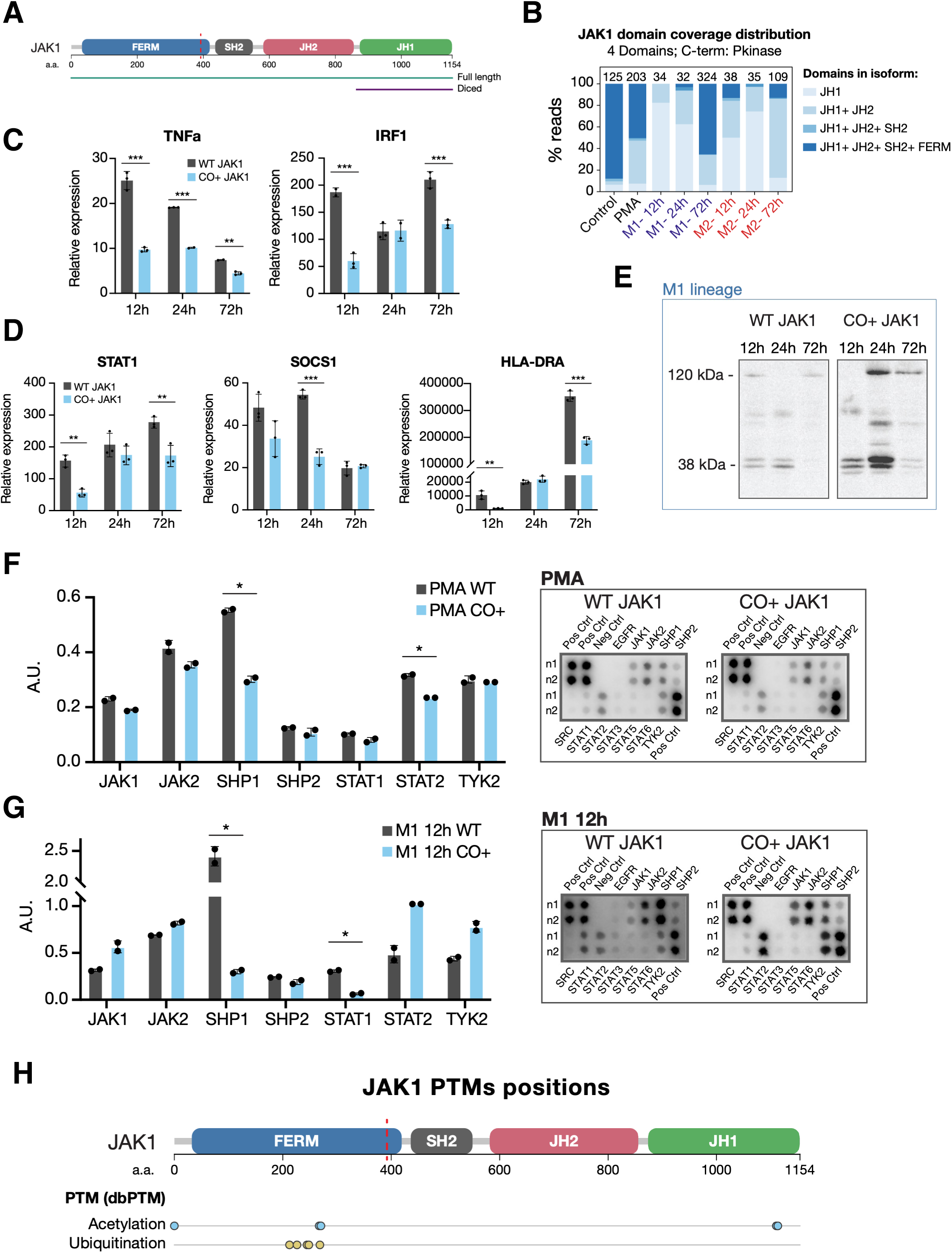
Dicing of JAK1 is required for M1 macrophage polarization. **(A)** Srtuctural ilustration of canonical full-length JAK1 and the diced JH1 kinase module. The dashed red line indicates the dicing site.**(B)** JAK1 RNA-dicing dynamics during M1 polarization determined by RNA-seq, showing maximal dicing during the early phase of polarization followed by a progressive return toward the PMA state by 72 h. **(C)** qRT-PCR analysis of the M1 markers TNFα and IRF1 in cells expressing wild-type JAK1 or the synonymous codon-modified, dicing-impaired JAK1 allele (CO+) at 12, 24, and 72 h following M1 induction. **(D)** Expression of the JAK1-associated genes STAT1, SOCS1, and HLA-DRA in wild-type and CO+ cells across M1 polarization. two-tailed Student’s t test; *\*p < 0.05, **p < 0.01 ***p < 0.001* **(E)** Representative JAK1-HA western blot analysis showing temporal expression of canonical and diced JAK1 isoforms in cells expressing wild-type or CO+ JAK1. Wild-type JAK1 undergoes a transient shift toward the diced JH1 isoform, with maximal JH1 abundance at 24 h, whereas synonymous recoding disrupts normal isoform timing and stoichiometry. **(F, G)** JAK/STAT pathway phosphorylation profiling in wild-type and CO+ cells after PMA treatment (F) and at 12 h following M1 induction (G). Wild-type cells preferentially engage a canonical JAK1-associated signaling state characterized by increased STAT1 and SHP1 phosphorylation during M1 induction. CO+ cells exhibit increased TYK2 and STAT2 phosphorylation. Unpaired *t*-test; *\*p < 0.05* **(H)** Positions of annotated JAK1 post-translational modifications mapped along the protein sequence. A cluster of ubiquitination sites at residues 213–269 lies within an N-terminal region retained in canonical JAK1 but absent from the diced JH1 module, providing a structural basis for differential post-translational regulation of the two isoforms.

We further showed that synonymous recoding of the JAK1 open reading frame substantially alters its dicing potential and produces marked biological effects, including changes in cellular proliferation. Therefore, we transduced HL-60 cells with vectors expressing HA-tagged JAK1 encoded either by the wild-type open reading frame or by a codon-modified allele in which approximately 20% of the nucleotides were altered through synonymous substitutions while preserving an identical amino acid sequence. These synonymous substitutions alter RNA-encoded determinants of dicing while preserving the canonical JAK1 protein sequence, thereby enabling selective perturbation of JAK1 dicing while minimizing contributions from other sources of transcript diversity, including alternative splicing and alternative promoter usage. RNA-seq analysis of JAK1 revealed a pronounced dicing signature during M1-like polarization, with maximal dicing at 12 h, followed by a progressive return toward the PMA state by 72 h (**Figure 4B; Figure S4A**). We therefore infected HL60 cells with vectors encoding either WT or codon-modified (CO+) JAK1, and induced M1-like differentiation as described. Following M1-like differentiation, the two cell types differed clearly in their morphology: wild-type JAK1 cells showed the differentiated clustering also seen in non-transfected HL-60 cells, whereas cells expressing the CO+ allele showed a weaker clustering phenotype (**Figure S4B**). Next, we performed qRT-PCR to assess the expression of the proinflammatory markers TNFα and IRF1, together with canonical downstream JAK1 targets, including SOCS3, STAT1, and HLA-DRA– dependent genes.

CO+ cells showed reduced expression of M1-like differentiation markers compared with WT JAK1 as early as 12 h, whereas M2-like differentiation markers were affected more modestly. (**Figure 4C; Figure S4C**), accompanied by reduced expression of downstream JAK1 targets (**Figure 4D**). We next performed western blotting for JAK1 using HA-tag antibody to differentiate our construct from endogenous JAK1, to track the dicing dynamics of each construct over time. Wild-type JAK1 expressed both canonical and diced JH1 isoforms at 12 h, shifted toward predominantly diced JH1 expression at 24 h, and returned to predominantly canonical expression by 72 h closely recapitulating the transient dicing dynamics observed at the RNA level (**Figure 4E**). In contrast, the CO+ JAK1 allele showed an unbalanced canonical-to-diced ratio at 12 h, elevated overall expression accompanied by additional intermediate isoforms at 24 h, and persistence of these intermediates through 72 h (**Figure 4C**). Because these synonymous changes are expected to disrupt the conserved, programmed dicing of JAK1, this dysregulated expression is consistent with the compensatory upregulation we reported previously upon the expression of dicing-deficient JAK1, resulting in altered timing and stoichiometry of canonical and diced isoform expression.

Next, we profiled JAK/STAT signaling using a phosphorylation ELISA array in cells expressing each construct after PMA treatment and 12 h after IFN + LPS exposure. Consistent with the qRT-PCR and RNA-seq data, JAK1 dicing was low before macrophage differentiation and was accompanied by limited downstream pathway activation. At 12 h after M1-like induction, cells expressing wild-type JAK1 showed increased phosphorylation of STAT1, a well-established JAK1 effector, together with SHP1, a negative regulator of JAK signaling (**Figure 4F–G**). In contrast, CO+ cells displayed increased TYK2 and STAT2 phosphorylation, indicating a shift away from the JAK1–STAT1 axis. Interestingly, such redistribution is consistent with the known JAK– STAT signaling cascade and the fact that perturbation of JAK1/TYK2 activity can reconfigure IL-10- and interferon-dependent feedback circuits ^24–26^, associated with macrophage states. Together, these findings provide functional evidence that disruption of JAK1 dicing alters pathway signaling during early macrophage differentiation.

Finally, the altered JAK1 isoform dynamics in CO+ cells relative to wild-type cells suggest that, beyond the regulation of isoform generation by dicing, canonical and diced JAK1 may be independently regulated at the post-translational level. Following the positional organization of PTM identified in **Figure 3G**, we asked whether PTM, which remains unaffected by synonymous recoding of the construct, differentially marks the canonical and diced JAK1 isoforms and potentially contributes to their distinct stability. Mapping of annotated PTM revealed a single cluster of ubiquitination sites at residues 213–269, positioned upstream and adjacent to the dicing site within the N-terminal region retained in canonical JAK1 but absent from the diced JH1 module. (**Figure 4H**). This physical organization shows that ubiquitination sites are distributed in relation to the dicing pattern, such that dicing partitions potential ubiquitination sites between the resulting isoforms, thereby providing a potential basis for independent post-translational regulation of the canonical and diced isoforms.

### RNA dicing mediates functionally distinct isoforms with divergent kinetic and signaling properties

We previously showed that the diced JH1 kinase domain and canonical full-length JAK1 differ in both phosphorylation activity and subcellular localization, leading to distinct biological outputs ^12^. The tightly regulated dynamics of JAK1 dicing shown in **Figure 4** further suggest that the JH1 module is not simply a truncated kinase that retains catalytic activity but a functionally distinct signaling entity. Consistent with this idea, previous studies have demonstrated substantial functional differences among alternative JAK domain architectures ^8,9,41^. Viewed in the context of RNA dicing, these observations suggest that the conventional linear JAK–STAT model represents an oversimplified view of JAK signaling. In contrast, modular, dicing-dependent protein architectures may diversify signaling outputs by altering substrate recognition, subcellular localization, and signaling dynamics ^42^.

Therefore, we hypothesized that RNA dicing expands kinome signaling diversity by generating modular kinase isoforms with distinct substrate specificities and signaling properties. To test this directly, we compared recombinant full-length JAK1 and the isolated JH1 kinase domain using a 14-point titration on a PamChip PTK peptide array containing 166 known tyrosine kinase substrates ^41^. Overall, we identified 79 peptides that responded to either JAK1 isoform. As predicted, the diced JH1 domain reached 80% of its maximal response (EC_80_) at a median of 57 ng of enzyme, compared with 542 ng for full-length JAK1, corresponding to a 9.5-fold lower EC (**Figure S5A**). Across the 20 shared substrates, the diced kinase consistently displayed greater potency (median 12-fold lower EC), which remained approximately 3.5-fold higher after normalization for molecular stoichiometry.

The increased potency of the diced JH1 domain alone could not distinguish between the enhanced catalytic activity and altered substrate response behavior. We therefore examined individual peptide dose-response relationships to determine whether the two JAK1 isoforms differed only in catalytic efficiency or also in the kinetics of substrate phosphorylation. Across the seventy-nine responsive peptides, the substrate repertoires proved to be largely non-overlapping rather than nested (**Figure 5A**). Thirty peptides were preferentially phosphorylated by full-length JAK1, whereas twenty-nine were preferentially phosphorylated by the diced JH1 kinase, with only 20 peptides robustly shared between the two isoforms. Importantly, these differences reflected changes in phosphorylation kinetics rather than catalytic potency alone. Full-length-preferred substrates exhibited classical sigmoidal dose-response curves with full-length JAK1 (median Hill slope 1.05; median R² = 0.95), but failed to generate interpretable dose-response relationships with the diced kinase, yielding poor fits and highly variable Hill coefficients, which are characteristic of substrates that do not enter a productive catalytic signature (**Figure S5B**). Conversely, JH1-preferred substrates remained phosphorylated by full-length JAK1 but displayed shallow, non-saturating dose-response curves (median Hill slope 0.59 versus 1.05 for the diced kinase), with many failing to reach their fitted plateau even at the highest enzyme concentration tested (**Figure S5B**). Thus, the RNA dicing machinery does not simply increase kinase activity but changes the phosphorylation pattern of individual substrates, enhancing the phosphorylation of JH1-preferred targets while weakening responses to alternative substrates. These findings reveal a potential functional consequence of RNA dicing, with increased catalytic potency and a shift in kinase substrate preference.

**Figure 5.**
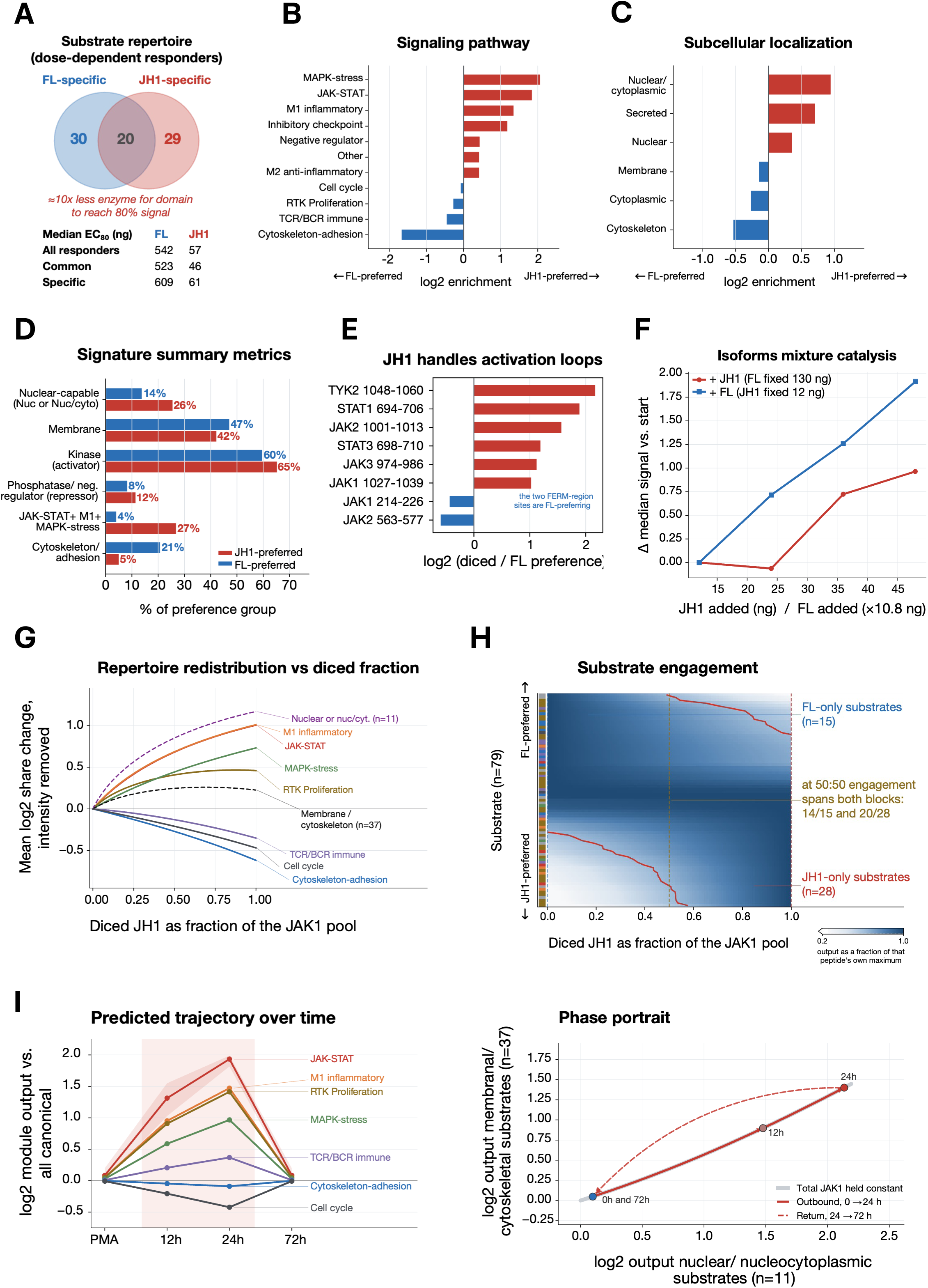
RNA dicing generates JAK1 isoforms with distinct substrate preferences and signaling outputs. **(A)** Substrate repertoires of full-length JAK1 and the diced JH1 kinase module determined by dose-response profiling on the PamChip PTK array. Of 79 responsive peptides, 30 preferentially responded to full-length JAK1, 29 to diced JH1, and 20 were shared. Median EC80 values illustrate the greater apparent potency of the diced kinase across responsive substrate classes. **(B–D)** Functional properties of substrates preferentially phosphorylated by diced JH1 or full-length JAK1. **(B)** Enrichment of biological and signaling categories among preferred substrates. **(C)** Subcellular localization of preferred substrates, showing enrichment of nuclear-associated targets for diced JH1 and membrane/cytoskeletal targets for full-length JAK1. **(D)** Summary of signaling features distinguishing the two substrate repertoires, including localization and pathway-associated preferences. **(E)** Relative phosphorylation preference of selected JAK-family and STAT peptides. Diced JH1 preferentially phosphorylates activation-loop peptides from multiple JAK-family members, whereas peptides from N-terminal JAK regulatory regions are preferentially phosphorylated by full-length JAK1. **(F)** Co-titration of full-length JAK1 and diced JH1, demonstrating that both kinase isoforms remain catalytically active in mixed reactions and contribute to the combined phosphorylation output. **(G)** Predicted redistribution of functional substrate classes as the diced JH1 fraction of the total JAK1 pool increases. Increasing JH1 shifts phosphorylation toward nuclear, JAK– STAT, M1-inflammatory, and stress-associated targets while reducing membrane, cytoskeletal, and adhesion-associated outputs. **(H)** Substrate engagement across increasing diced-JH1 fractions, illustrating the progressive recruitment and loss of distinct substrate groups as JAK1 isoform stoichiometry changes. **(I)** Predicted temporal trajectory of pathway-level signaling output obtained by integrating the biochemical substrate profiles with endogenous JAK1 isoform dynamics during M1 polarization presented in Figure 4. **(J)** Phase-space representation of predicted nuclear versus membrane/cytoskeletal signaling output, illustrating the transient signaling excursion during early M1 polarization and return toward the initial state as JAK1 dicing resolves.

To determine whether these biochemical differences correspond to distinct biological functions, we grouped the preferential substrates according to the subcellular localization and functional annotations of their corresponding proteins. For pathway enrichment, all peptides exhibiting directional preference were included. The diced JH1 domain preferentially targeted nuclear-associated proteins together with core JAK– STAT, M1-like inflammation-associated, stress MAPK, and immune receptor signaling pathways. In contrast, full-length JAK1 preferentially phosphorylated proteins involved in membrane organization, cytoskeletal regulation, and cell adhesion (**Figure 5B–D**, **Figure S5C**). The enrichment of M1-like inflammatory substrates closely matched the requirement for JAK1 dicing during proinflammatory macrophage differentiation (**Figure 4**), while the enrichment of nuclear-associated substrates was consistent with the loss of the FERM membrane-targeting domain and nuclear re-localization of the JH1 domain we previously observed ^12^. Interestingly, the diced kinase preferentially phosphorylated activation-loop peptides from multiple JAK family members, including the JAK1 activation loop (Y1034/Y1035), whereas peptides derived from the N-terminal regulatory region (residues 214–226) were preferentially phosphorylated by full-length JAK1 (**Figure 5E**). Strikingly, this N-terminal regulatory region overlaps with the ubiquitination sites mapped in **Figure 4H**, indicating that the canonical isoform contains a regulatory module subject to multiple post-translational inputs, including ubiquitination and auto-phosphorylation. These observations provide a mechanistic basis for the notion that increased JH1 abundance may promote trans-phosphorylation of full-length JAK1 while simultaneously shifting signaling toward the nucleus. Together, these findings demonstrate that RNA dicing rewires the intrinsic kinase substrate preference in a manner that predicts the distinct cellular functions of the two JAK1 isoforms.

As both kinase isoforms are generated from the same canonical transcript and coexist during the dicing process, progressive RNA dicing is predicted to alter not only substrate phosphorylation but also the relative predicted abundance of full-length homodimers, full-length JH1 heterodimers, and JH1 homodimers. We therefore asked how progressive changes in isoform stoichiometry affect the phosphorylation landscape. To address this, we profiled seven defined mixtures of recombinant full-length JAK1 and diced JH1 on the same PamChip PTK array, spanning 6.5–52.6% diced/full-length fractions on a molar basis.

Both isoforms remained catalytically active throughout the mixture, and neither inhibited the activity of the other. Instead, the array response was well described by a simple additive two-component model, indicating that each isoform contributes independently to the overall phosphorylation pattern (**Figure 5F**). Increasing the abundance of either isoform selectively enhances the phosphorylation of its preferred substrates. Although the diced JH1 domain appeared more active on a mass basis, this difference was largely explained by its smaller molecular weight, with both isoforms displaying similar molar catalytic activities. These findings suggest that the two isoforms function cooperatively rather than independently. As the full-length: JH1 ratio changes, phosphorylation is progressively redistributed between their complementary substrate repertoires, whereas the diced JH1 isoform may simultaneously promote the activation of full-length JAK1 through trans-phosphorylation of its activation loop. Although overall kinase activity remained largely unchanged, altering the relative abundance of full-length JAK1 and diced JH1 profoundly reshaped substrate phosphorylation. Rather than generating new signaling activities, progressive RNA dicing redistributed phosphorylation between complementary substrate repertoires, thereby continuously remodeling the signaling output as the isoform stoichiometry changed (**Figure 5G-H**).

Finally, we integrated these biochemical measurements with the endogenous dynamics of JAK1 dicing during macrophage polarization (**Figure 4**). The transient increase in the diced JH1 isoform predicts a coordinated redistribution of signaling toward nuclear-associated proteins, core JAK–STAT signaling, M1-like inflammatory pathways, stress MAPK signaling, and immune receptor signaling, while reducing the phosphorylation of membrane-associated, cytoskeletal, adhesion, and cell cycle-associated targets (**Figure 5B–D**; **Figure S5D-E**). Simultaneously, the increasing proportion of JH1-containing complexes is predicted to favor activation-loop phosphorylation and nuclear accumulation of the diced kinase, thereby coupling changes in dimer composition, subcellular localization, and substrate selection into a coordinated signaling transition (**Figures 5G-E**). Collectively, these observations indicate that transient RNA dicing progressively redirects signaling toward the nuclear JAK–STAT, inflammatory, and stress MAPK pathways while reducing membrane-associated and cytoskeletal signaling. Thus, dynamic changes in JAK1 isoform stoichiometry are translated into coordinated remodeling of downstream signaling networks. Overall, these findings establish RNA dicing for JAK1 as a mechanism that converts a single kinase into a dynamic signaling system by remodeling substrate phosphorylation kinetics. Progressive changes in protein modular architecture reshape substrate accessibility, pathway selection, and downstream signaling output, providing a mechanistic basis for the functional diversification of kinase signaling from a single genetic locus.

## Discussion

In this study, we define RNA dicing as a global, robust, and adaptive layer of post-transcriptional gene regulation. Rather than representing constitutive RNA turnover, RNA dicing operates as a transient program of transcriptome remodeling that is preferentially engaged during the early stages of macrophage polarization before stable transcriptional programs are established. By remodeling the coding architecture of mature transcripts, dicing generates independently translated RNA modules, thereby expanding the gene output without requiring de novo transcription. Long-read sequencing further demonstrated that dicing converts intact transcripts into distinct coding molecules and preferentially occurs at positions related to protein domain architecture.

A defining feature of RNA dicing is its temporal organization. Dicing is most prominent during the early phase of macrophage polarization and progressively subsides as stable transcriptional programs emerge. This temporal separation suggests that transcription and RNA dicing serve complementary roles during cell-state transitions. While anabolic transcription establishes a longer-term cellular identity through the synthesis of new RNA, catabolic RNA dicing can rapidly alter gene output by remodeling pre-existing canonical transcripts. In this framework, the transcriptome defines a repertoire of potential protein outputs, whereas dicing determines which outputs are transiently deployed in response to changing cellular demands. Such regulation provides a mechanism for rapid but reversible adaptation during transitional cell states. From an immunological perspective, this model offers a clear functional advantage by enabling rapid and adjustable adaptation to changing immune demands, while minimizing the energetic cost associated with de novo gene expression.

Our findings further suggest that RNA dicing alters not only transcript architecture but also the organization of cellular signaling. Canonical models of signaling diversity generally assume that a gene produces a principal protein whose activity is subsequently diversified through post-translational regulation. RNA dicing adds an earlier layer of diversification by generating protein isoforms with distinct domain compositions, catalytic properties, substrate preferences, localizations, and signaling behaviors. The dynamic interaction between canonical JAK1 and its diced JH1 module illustrates this principle. In addition to the canonical functions of full-length JAK1, the diced JH1 isoform displays enhanced catalytic activity, distinct activation kinetics, preferential nuclear localization, and a signaling repertoire essential for M1-like proinflammatory macrophage polarization. Thus, dicing can expand the signaling potential of a single gene by generating functionally distinct protein architectures from common transcripts.

This principle may extend well beyond JAK1. Decades of deletion and domain-mapping experiments have shown that isolated protein regions and truncated proteins can display activities that differ markedly from, and in some cases oppose, those of their full-length counterparts, including dominant-negative behavior ^43^. These observations have established the functional autonomy of many protein domains but often lack a physiological mechanism explaining how analogous products might arise endogenously. Our findings suggest that RNA dicing may provide such a mechanism by selectively generating protein isoforms with altered domain composition. In this view, protein domains are not only structural components of full-length proteins but also potential functional modules that can be differentially deployed through regulated processing of mature RNAs.

The relationship between RNA dicing and protein-domain organization also has consequences for post-translational regulation. The removal of defined coding regions necessarily alters the repertoire of regulatory sites retained within the resulting protein isoforms, including phosphorylation, ubiquitination, acetylation, and other modifications. Therefore, dicing does not need to directly regulate post-translational modifications to alter their functional impact; by physically removing specific protein domains, it determines which regulatory elements and modification sites remain available in each resulting isoform. The spatial correspondence observed between JAK1 dicing, regulatory domains, and post-translational modification sites supports this modular organization and suggests that distinct diced isoforms may acquire different stabilities and regulatory properties after translation.

The temporal and lineage-specific nature of dicing further indicates that this process is not simply an intrinsic property of individual transcripts. Differential dicing in pro-inflammatory M1-like and anti-inflammatory M2-like macrophages shows that the same transcript can be remodeled differently depending on cellular context, producing distinct functional outputs adapted to lineage-specific demands. This observation implies the existence of trans-acting mechanisms that regulate transcript selection, cleavage position, or both. Consistent with this idea, STAT1 and STAT3 undergo extensive but distinct remodeling during M1- and M2-like polarization, further linking dicing to cell-state-specific signaling programs.

More broadly, RNA dicing provides a framework for understanding why measurements of total transcript abundance often correlate only incompletely with protein abundance or biological phenotypes ^1,2^. Conventional transcriptomic approaches typically treat each gene as a single quantitative unit, yet our findings indicate that transcripts from the same locus can occupy distinct functional states depending on how their coding architecture has been remodeled. Consequently, the biological output may change substantially, even when the total gene expression changes only modestly. Resolving transcript architecture at the intra-gene level may therefore reveal regulatory information that is obscured by conventional gene-level measurements and help connect RNA abundance more directly to proteomic and phenotypic outcomes.

From an evolutionary perspective, RNA dicing offers an efficient means of expanding functional diversity through the reuse of existing coding information. Rather than requiring a dedicated gene for every functional context, cells can potentially generate different combinations of conserved protein modules from the existing transcripts. During rapid cellular transitions, such mechanisms could provide an immediate means of altering protein output before slower transcriptional programs are fully established. Therefore, the differential deployment of existing RNA and protein modules may represent an economical strategy for increasing regulatory flexibility without the proportional expansion of genome size. More generally, RNA dicing may represent one example of a broader evolutionary principle in which biological complexity emerges through modular organization and combinatorial reuse of existing components.

However, several important mechanistic questions remain. Although our previous work established APA-directed cleavage and m6A-dependent translation as core features of RNA dicing, additional mechanisms are likely to contribute to transcript recognition and cleavage ^10–12^. Preliminary observations suggest that sequence-specific endonucleases and RNA sequence- or structure-dependent targeting mechanisms may converge on similar modular outcomes; however, these possibilities require direct experimental testing. The determinants that specify lineage- and state-dependent dicing also remain unknown. Likewise, the relationship between dicing and post-translational regulation warrants further investigation, particularly whether isoform-specific loss or retention of regulatory sites systematically alters protein stability, localization and signaling activity.

Taken together, our findings support a model in which the mature transcriptome remains functionally programmable, even after transcription. Transcription establishes the repertoire of available gene products, whereas RNA dicing dynamically determines which protein modules are generated and deployed during cellular adaptation. In this framework, gene function cannot always be inferred from transcript abundance or canonical protein sequences alone, but also depends on the evolving architecture of mature RNAs and the protein isoforms they produce. Therefore, RNA dicing provides a mechanistic link between transcript architecture, proteomic diversity, signaling behavior, and cellular phenotype at the intra-gene resolution. If this principle extends broadly across tissues and developmental contexts, it may require a more modular view of how gene regulation and genotype–phenotype relationships are interpreted.

## Lead contact

Further information and requests for reagents and resources should be directed to the lead contact Yuval Malka.

## Materials availability

Plasmids generated in this study are available from the lead contact upon reasonable request and are subject to institutional material transfer requirements.

## Acknowledgment

Kinome profiling experiments were supported in part by funding from the Biotech Booster program through project BB25034 (DUS-I reference BIOB25006).

## Authors Contribution

Y.M. conceived and supervised the project, designed experiments, analyzed the data, and wrote the manuscript. N.T. and O.Y. designed and performed the experiments and analyzed the data. D.H.Y., T.B., and H.T. analyzed the data. R.A.-R. performed experiments.

## Data availability

The long-read RNA-sequencing data analyzed in this study were generated by ENCODE and are available through the Gene Expression Omnibus under the accession number GSE219923. The mass spectrometry data analyzed in this study were generated in our previous work and are available through ProteomeXchange under accession PXD071451. The publicly available m6A-RIP-seq data analyzed in this study are available through the Gene Expression Omnibus under the accession number GSE112795.

## Materials and Methods

### Experimental model and cell culture

Human promyelocytic HL-60 cells were maintained at 37°C in a humidified 5% CO2 atmosphere in Dulbecco’s modified Eagle’s medium supplemented with 20% fetal calf serum, 100 U/mL penicillin, and 100 μg/mL streptomycin. Cells were differentiated into macrophage-like cells with 10 μM phorbol 12-myristate 13-acetate (PMA) for 120 h ^44^. PMA-differentiated cells were polarized toward an M1-like state with 100 ng/mL lipopolysaccharide (LPS) and 100 ng/mL interferon-γ (IFN-γ), or toward an M2a-like state with 10 ng/mL interleukin-4 (IL-4) and 10 ng/mL interleukin-13 (IL-13), for 12, 24, or 72 h.

### Public long-read RNA-seq data and transcript-integrity analysis

Long-read RNA-seq datasets were obtained from the ENCODE portal ^27^. The analysis included untreated HL-60 cells (ENCSR887LTD), PMA-treated HL-60 cells at 120 h (ENCSR546DFO), M1-polarized HL-60 cells at 12, 24, and 72 h (ENCSR121FDE, ENCSR583KAF, and ENCSR398SKD, respectively), and M2a-polarized HL-60 cells at 12, 24, and 72 h (ENCSR278ZPI, ENCSR930GRQ, and ENCSR159ICU, respectively). These datasets were generated in the Mortazavi laboratory.

The reads were classified relative to the annotated canonical coding sequence (CDS). Reads containing the canonical translation initiation site but lacking the annotated in-frame stop codon were classified as 5′ transcripts. Reads containing complete canonical CDS were classified as full-length (FL) transcripts. Reads that retained the canonical in-frame stop codon but lacked the canonical translation initiation site were classified as 3′ transcripts. The 3′ class was used as an operational signature for downstream RNA-dicing products.

### Differential Gene Expression Analysis (DESeq2)

Gene-level count tables were generated by summing the transcript reads per gene from the sample files, excluding TALON-specific internal identifiers (ENCL). Low-abundance genes detected in fewer than three samples or with fewer than 10 total reads summed across all samples were removed prior to analysis. Differential expression was evaluated using the **DESeq2** package in R with the generalized linear model design ∼ group, using unpolarized cells (control) as the reference level. Gene annotations were retrieved using the **biomaRt** package. Pairwise differential testing (Wald test) was conducted across baseline controls (vs. Control and PMA), time-matched polarization lineages (M1/IFN vs. M2/IL at 12, 24, and 72 h), and consecutive time-point transitions within each lineage. P-values were corrected for multiple testing using the Benjamini-Hochberg False Discovery Rate (FDR) method at α=0.05. Genes with p_adj_ ≤ 0.05 and | log2 Fold Change ≥1 were designated as significantly differentially expressed.

### Differential Transcript Truncation and Isoform Composition Analysis (DRIMSeq)

To evaluate post-transcriptional dicing dynamics independently of total gene abundance, transcript architecture was classified into three terminal read categories (start_only, stop_only, and both) for each gene using **DRIMSeq software**. Datasets were filtered to retain features with ≥5 counts and genes with ≥10 total counts in at least two samples (dmFilter).

For each pairwise comparison (matching the DESeq2 contrasts), a Dirichlet multinomial model was fitted using the design formula ∼ group. Dispersion and gene-wise precision were calculated (dmPrecision), followed by proportion estimation (dmFit) and Likelihood Ratio Testing (dmTest). Significance was corrected using the Benjamini-Hochberg procedure, defining dicing/truncation events at p_adj_ ≤0.05. Global and gene-specific relative proportions across feature categories were extracted to quantify the structural shifts across the polarization stages.

### Protein-domain and transcript-boundary analyses

Protein domain coordinates were obtained from Pfam-A annotations projected onto the human genome (hg38), as distributed through the UCSC Genome Browser “Pfam in UCSC Genes” track. Domains were mapped to transcript coding coordinates using UCSC knownGene models (hg38), with genomic boundaries converted in a strand-aware manner to CDS nucleotide and corresponding amino-acid positions. For each downstream 3′ transcript, the neo-5′ boundary was defined as the first aligned nucleotide and classified as either within an annotated protein domain or within an inter-domain coding region.

To test whether neo-5′ boundaries were non-randomly positioned relative to protein domains, the observed distribution was compared with a length-proportional uniform null model in which every coding residue had an equal probability of serving as a neo-5′ boundary. Expected frequencies within domains versus inter-domain regions, or upstream versus downstream of domain starts, were therefore proportional to the number of coding residues available in each category. Departure from this expectation was assessed using a binomial test at the site level and independently confirmed across proteins using the Wilcoxon signed-rank test.

Domain coverage was determined by identifying all the annotated protein domains retained downstream of each neo-5′ boundary. For multidomain proteins, the retained domains were counted from the C-terminus to quantify the preservation of downstream modular architectures.

### Pathway enrichment analysis

Genes identified by differential transcript usage or gene-level differential expression were evaluated for enrichment in the MSigDB Hallmark gene sets ^45^. Analyses were performed separately for the M1 and M2 comparisons and time-resolved transitions.

### m6A-site preparation and positional analysis

FASTQ reads were aligned upstream of this workflow to the Homo_sapiens.GRCh38.dna.primary assembly.fa reference genome using HISAT2 ^46^. This pipeline began with the resulting BAM files.

### CDS filtering

A temporary GTF containing only CDS features was generated from the reference annotation. BAM files were intersected with the CDS-only annotation using bedtools intersect -f 1.0, retaining reads fully contained within a CDS interval and excluding reads with partial overlap or alignment extending into intronic or untranslated regions (UTRs). The filtered BAM files were indexed using samtools.

### Transcript-level read assignment

The GTF was parsed to generate transcript-specific exon coordinates, CDS start and end positions, and total CDS lengths. Transcript records were indexed by chromosome using an interval tree. Each read was assigned only to candidate transcripts that overlapped with their genomic span, shared their strand, and contained every alignment block entirely within a CDS exon of the same transcript. This transcript-level validation was required because the preceding CDS filter was reference-wide, rather than transcript-specific.

### m6A-site preparation

m6A-IP-seq data were obtained from the GEO accession GSE213207. The m6A coordinates were derived from Pinello et al.. CDS intervals were merged separately by strand to generate a ^47^erence coding track. m6A sites located within 100 nt of the start or end of the overlapping merged CDS interval were excluded.

Independently of this boundary filter, every m6A site overlapped with raw transcript-level CDS records to generate a permissive list of compatible transcript IDs and gene names. The final transcript-level assignment required both the m6A site and the read-defined transcript boundary to occur within the CDS exons of the same transcript. The genomic sequence underlying each m6A site was extracted from the reference FASTA file for quality control. The distances between m6A positions and long-read transcript 5′ boundaries were calculated in spliced transcript coordinates and compared with two transcript-matched null models: a fixed transcript-midpoint control and a shuffled control, in which for each observed read–m6A pair, a random position was selected from the same transcript. These controls preserved transcript identity while removing the observed positional relationship between the neo-5′ boundary and m6A. Analyses were restricted to m6A sites located ≥100 nt from the CDS boundaries, and distance distributions were visualized within ±500 nt of the m6A-site center. Replicates were pooled by condition to compare the observed and control distributions.

### RNA isolation, reverse transcription, and qPCR

Total RNA was isolated using the Quick-RNA Miniprep Kit (Zymo Research, R1055) according to the manufacturer instructions. RNA concentration was measured using a NanoDrop spectrophotometer. cDNA was generated using the LunaScript RT Master Mix Kit (New England Biolabs, E3025) with Oligo d(T)23 VN primer (New England Biolabs, S1327S), following the manufacturer’s instructions. Quantitative PCR was performed using Fast SYBR Green Master Mix (Applied Biosystems, 4385614) following the manufacturer’s instructions.

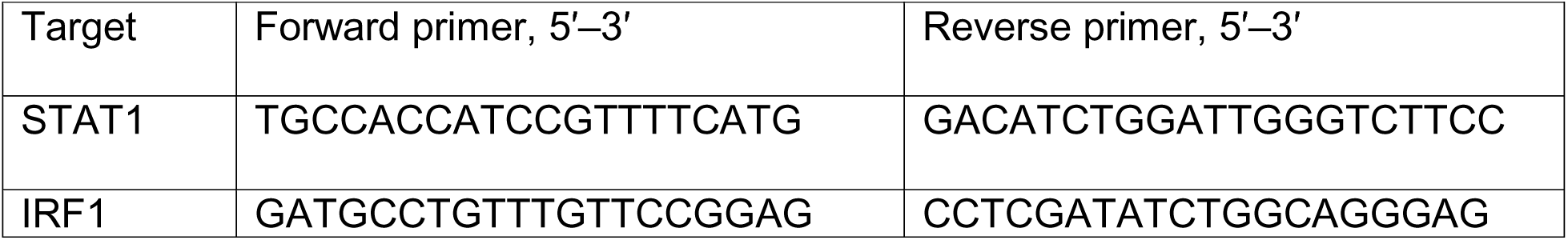
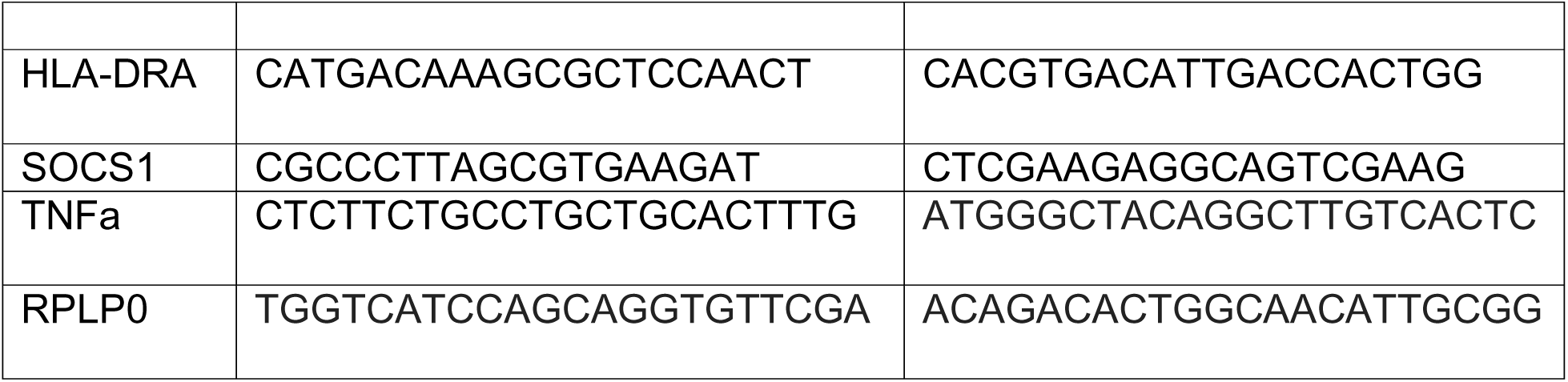

### Size-resolved proteomics

Size-resolved proteomics was performed as described by van der Kammen et al. (2026). Proteins were separated by SDS–PAGE and excised as light (10–40 kDa), intermediate (40–80 kDa), and heavy (80–180 kDa) fractions before mass spectrometry analysis. Peptide intensities were summarized at the gene level within each fraction. For proteins with canonical molecular masses ≥80 kDa, redistribution from the heavy to light fraction was used as evidence for lower-molecular-weight protein products. Proteins with a canonical molecular mass ≤40 kDa served as the fractionation control.

Light-to-heavy and heavy-to-light abundance ratios were calculated for each gene and compared between genes identified by both DESeq2 and DRIMSeq and those uniquely identified by DESeq2.

### JAK1 constructs and transduction

HA-tagged wild-type and synonymous codon-modified JAK1 constructs were generated, packaged, and transduced as previously described (van der Kammen et al., 2026). The codon-modified allele contains synonymous substitutions across approximately 20% of the JAK1 coding sequence while preserving the canonical amino acid sequence. Cells expressing wild-type or codon-modified JAK1 were differentiated with PMA and polarized toward M1 macrophages, as described above.

### Western blotting

Western blotting was performed as described by van der Kammen et al. (2026). Proteins were resolved on 4–15% gradient TGX gels (Bio-Rad) using the Laemmli buffer system and transferred to 0.2-μm nitrocellulose membranes (Pall). The membranes were blocked with 5% nonfat dried milk in TBST. Primary antibodies were used at the following, HA tag (clone EPR22819-101; Abcam, ab236632; 1:1,000) secondary antibodies were used at 1:10,000.

### JAK–STAT phosphorylation profiling

JAK–STAT phosphorylation was profiled using the RayBiotech AAH-JAKSTAT-1 phosphorylation array in wild-type and codon-modified JAK1 cells after PMA differentiation and at 12 h after M1 induction, using biological duplicates. The arrays were processed according to the manufacturer’s instructions. Signals were background-corrected and normalized, and downstream analyses were performed according to the manufacturer’s recommended workflow.

### JAK1 kinase peptide-array profiling and dose–response analysis

Recombinant full-length human JAK1 and isolated catalytic JH1 domain (residues 862– 1154) were produced in Sf9 insect cells using pFastBac constructs, as described by van der Kammen et al. (2026). Proteins were purified using Ni2+-affinity chromatography, followed by size-exclusion chromatography. The JH1-domain preparation was purified using 10% glycerol in all buffers.

Protein tyrosine kinase activity was profiled using KinomePro/PamDx on PamGene PamStation peptide-tyrosine-kinase arrays. The assay was performed using the dynamic PamChip format described by Sanz et al. (2011). The arrays were blocked with 2% BSA and washed with kinase buffer containing 50 mM Tris-HCl (pH 7.5), 10 mM MgCl2, 1 mM EGTA, 2 mM DTT, and 0.01% Brij-35. Kinase reactions were performed at 30°C in a final volume of 25 μL with 100 μM ATP and fluorescein-conjugated PY20 anti-phosphotyrosine antibody (12.5 μg/mL). The reactions were pumped through the porous array matrix for 60 cycles at 2 cycles/min and imaged every second cycle with integrated CCD detection.

Full-length JAK1 and JH1 were each tested at 14 enzyme inputs: 0.064, 0.32, 1.6, 8, 40, 102.4, 200, 256, 640, 1,000, 1,600, 4,000, 5,000, and 10,000 ng. These inputs corresponded to approximately 0.53 fmol–83.33 pmol for full-length JAK1 and 1.6 fmol– 250 pmol for JH1. Peptide phosphorylation was quantified as background-corrected log2 spot intensity on a 0–16 scale.

For each peptide and isoform, the vendor pipeline fitted a four-parameter logistic model in log10 enzyme input: where is the enzyme input in grams, Bottom and Top are the fitted asymptotes, logEC50 is the log10 half-maximal enzyme input, and is the Hill slope. The pipeline reports EC50, EC80, logEC80, Hill slope, dynamic range, and coefficient of determination. Among the 166 peptides with at least one fitted curve, 166 full-length-JAK1 and 162 JH1 fits were returned.

Peptides were classified as dose-responsive when and. This identified 50 full-length-JAK1-responsive and 49 JH1-responsive peptides. The shared peptides that met both criteria for both isoforms were included (n = 20). Full-length-preferred peptides passed the criteria for full-length JAK1 but not JH1 (n = 30), and JH1-preferred peptides passed the criteria for JH1 but not full-length JAK1 (n = 29). The remaining 67 peptides were not classified. “Preferred” therefore denotes an interpretable dose-dependent response under the stated criteria; it does not necessarily mean that the other isoform had no activity.

The median EC80 values based on the protein mass input were 542.39 ng for all full-length-JAK1-responsive peptides and 57.20 ng for all JH1-responsive peptides. For the shared peptides, the median EC80 values were 522.88 ng for full-length JAK1 and 46.41 ng for JH1. The median EC80 values for the full-length-preferred and JH1-preferred sets were 609.30 ng and 60.88 ng, respectively. These values are reported as mass-based apparent potency because a comparison of intrinsic catalytic efficiency requires normalization to active enzyme molarity and active protein fraction.

The fractional saturation at the enzyme input was calculated as

The working doses were each isoform’s EC80:4.33 pmol (520 ng) for full-length JAK1 and 1.2 pmol (48 ng) for JH1. In Supplementary Figure 5, fits with are shown as solid curves and lower-quality fits as dashed curves. Fits used for local saturation correction in fixed-dose analyses were required to meet and Confidence intervals for peptide populations were calculated using bootstrap resampling of peptides (20,000 resamples; fixed random seeds).

### Kinase-array substrate annotation and enrichment

Peptides were assigned to cognate proteins using UniProt accessions supplied with PamGene/KinomePro peptide-array annotation. Proteins were assigned to predefined pathway and subcellular localization categories, including nuclear, membrane/cytoskeleton/adhesion, JAK–STAT, M1-inflammatory, stress-MAPK, and immune-receptor signaling, using a manually curated scheme built from Gene Ontology (Cellular Component terms for subcellular localization and Biological Process terms for pathway assignment), UniProt/Swiss-Prot subcellular location annotations, and Reactome pathway memberships [insert the releases/versions and access date you actually queried, for example “GO release 2026-05-01, UniProt release 2026_02, Reactome v88, accessed May 2026”]. Each protein was assigned to a single dominant category by the following curation rules: a protein was placed in a signaling category (JAK–STAT, M1-inflammatory, stress-MAPK, immune-receptor) when annotated to the corresponding pathway term; proteins not mapping to any signaling pathway were assigned to a localization category (nuclear, or membrane/cytoskeleton/adhesion) on the basis of their primary annotated compartment; ties or multi-category proteins were resolved by [state your rule, for example, “the most specific term” or “consensus of two independent curators, discordances adjudicated by a third”]. Enrichment of each category among full-length-JAK1-preferred and JH1-preferred peptide sets was tested against all 79 dose-responsive peptides using a one-sided Fisher’s exact test on 2×2 contingency tables (category membership × peptide-set membership), evaluating the over-representation of each category in each preferred set relative to the full 79-peptide background. P-values were adjusted using the Benjamini–Hochberg procedure, and an adjusted P < 0.05 was considered significant.

### Post-translational-modification and protein-domain analysis

Experimentally supported PTM sites in human proteins were retrieved from dbPTM ^48^. Each site was mapped to its corresponding canonical UniProt sequence, and its relative position was defined as the modified residue number divided by the canonical protein length (0 = N-terminus; 1 = C-terminus).

Functional protein groups were defined by the Pfam membership ^49^. For each of the 12 groups, human proteins (UniProt taxonomy ID: 9606) containing at least one of a predefined set of group-defining Pfam accessions were retrieved using the UniProt REST API (e.g., PF00069 for protein kinases). Protein lengths and domain-boundary coordinates were obtained from the corresponding UniProt canonical sequence feature annotations.

For visualization, the relative positional distributions of acetylation and ubiquitination sites were estimated using Gaussian kernel-density estimation with Scott’s-rule bandwidth (SciPy v1.12.0). Statistical analyses were restricted to proteins containing at least one acetylation and one ubiquitination site. For each protein, a PTM-polarity offset was calculated as the mean relative position of its ubiquitination sites minus the mean relative position of its acetylation sites, such that positive values indicated a more C-terminal distribution of ubiquitination relative to acetylation. Within each functional group, the mean offset was tested against zero using a two-sided one-sample t-test and independently confirmed using the Wilcoxon signed-rank test. P values were Bonferroni-corrected across the 12 groups, and the effect size was reported as Cohen’s d, calculated as the mean offset divided by its standard deviation across proteins. JAK1 ubiquitination sites were mapped to the canonical JAK1 sequence (UniProt P23458) and evaluated relative to the RNA-dicing boundary.

Domain coverage was quantified by identifying all annotated Pfam domains retained downstream of each neo-5′ boundary. For multidomain proteins, retained domains were counted from the C-terminus to assess the preservation of the downstream modular architecture.

### Quantification and statistical analysis

Dose–response analyses of PamGene/KinomePro array data used vendor-supplied four-parameter logistic fits and the responder thresholds described above. Because each enzyme–peptide titration did not include independent replicate arrays, the individual peptide-level parameters were not interpreted in isolation. The per-peptide noise floors, calculated from the differences between independent DMSO arrays, were 0.249 log2 units for full-length JAK1, 0.330 for JH1, and 0.215 for the mixed-isoform condition. The uncertainty for peptide populations was estimated by bootstrap resampling of peptides rather than treating peptides as biological replicates.

For qRT–PCR, western blotting, size-resolved proteomics, and other cell-based assays, the number of independent biological replicates, statistical tests, sidedness, error-bar definitions, and significance thresholds are specified in the relevant figure legends and source data files. Unless otherwise indicated, the data are presented as mean ± SD. For differential gene expression, transcript usage, and enrichment analyses, P values were adjusted using the Benjamini–Hochberg procedure, and adjusted P < 0.05 was considered significant. For PTM polarity analyses, the P values were Bonferroni-corrected across the 12 functional groups.

## Notes

### Competing Interest Statement

The authors have declared no competing interest.

### Summary of Updates

This is updated version of the Co-authors

