## Supplementary Figures for "Transient RNA dicing reprograms functional transcriptome architecture during macrophage polarization": Supplementry.pdf

#### Supplementary Figure 1

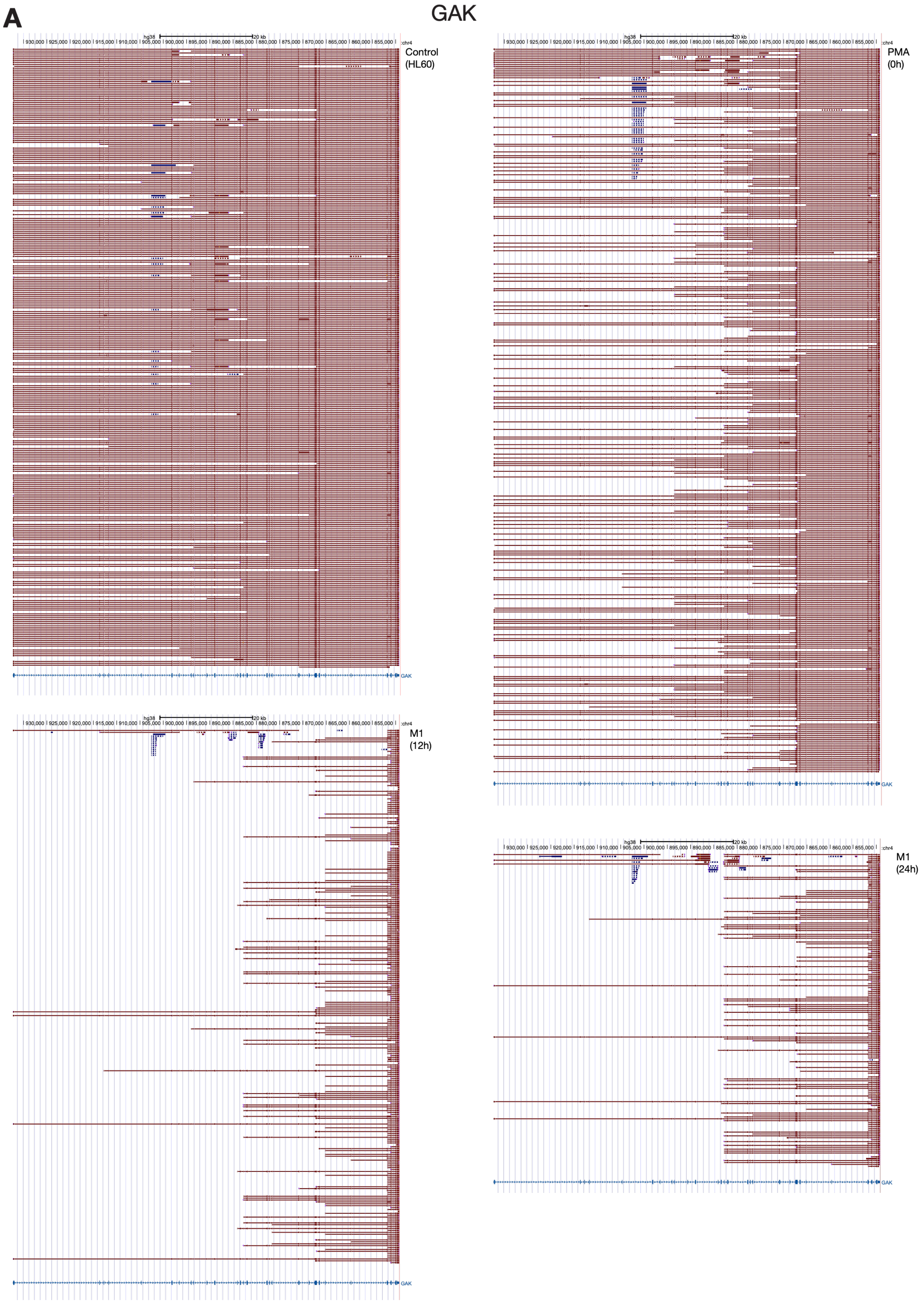

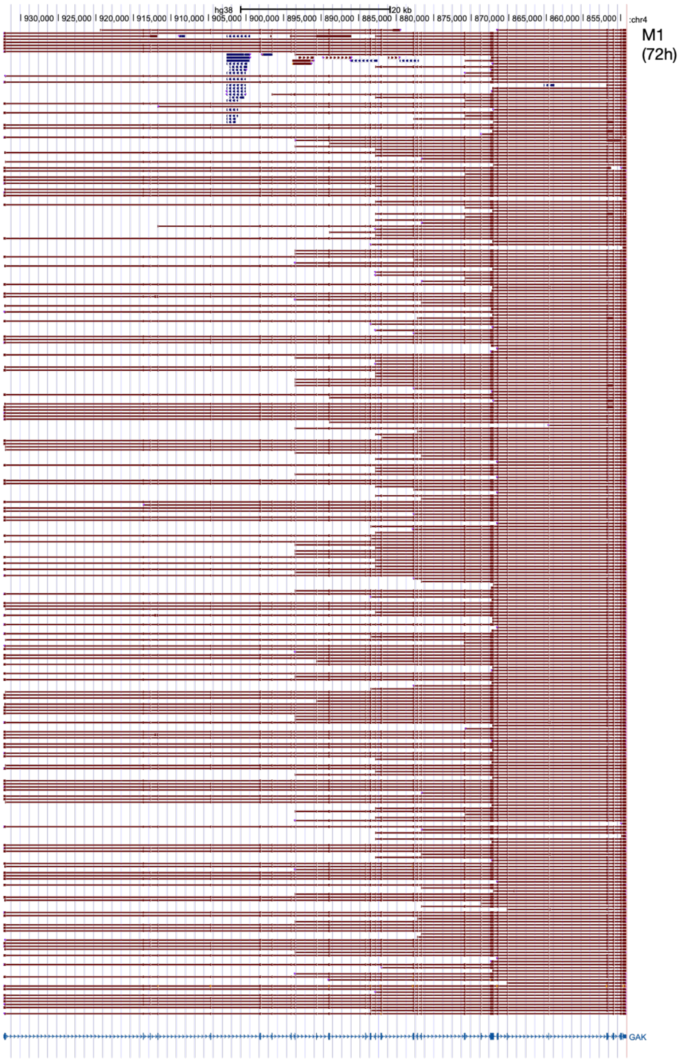
hg38
20 kb
930,000
925,000
920,000
915,000
910,000
905,000
900,000
895,000
890,000
885,000
880,000
875,000
870,000
865,000
860,000
855,000
:chr4
M1
(72h)
GAK

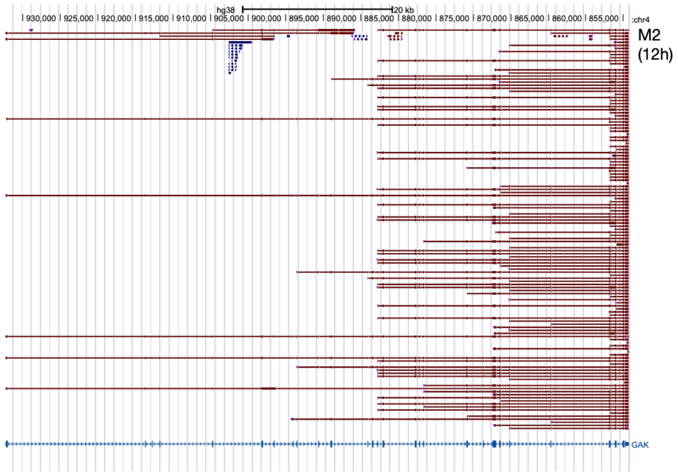
hg38
20 kb
930,000
925,000
920,000
915,000
910,000
905,000
900,000
895,000
890,000
885,000
880,000
875,000
870,000
865,000
860,000
855,000
:chr4
M2
(12h)
GAK

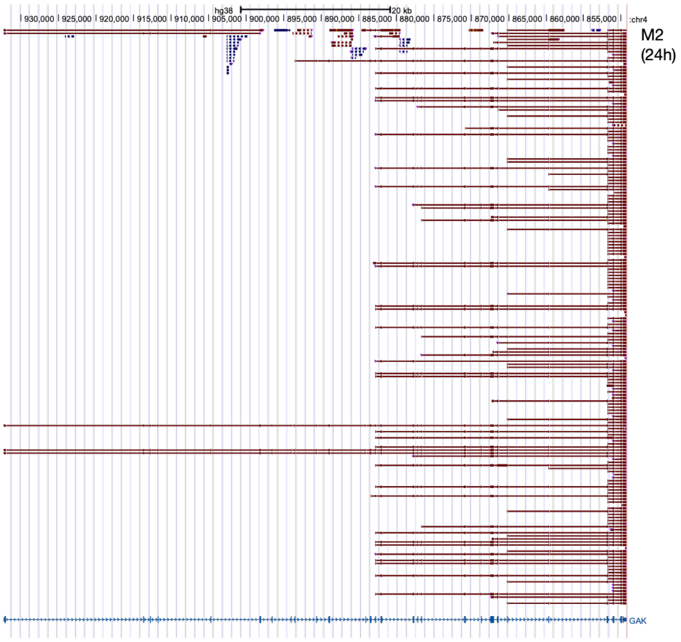
hg38
20 kb
930,000
925,000
920,000
915,000
910,000
905,000
900,000
895,000
890,000
885,000
880,000
875,000
870,000
865,000
860,000
855,000
:chr4
M2
(24h)
GAK

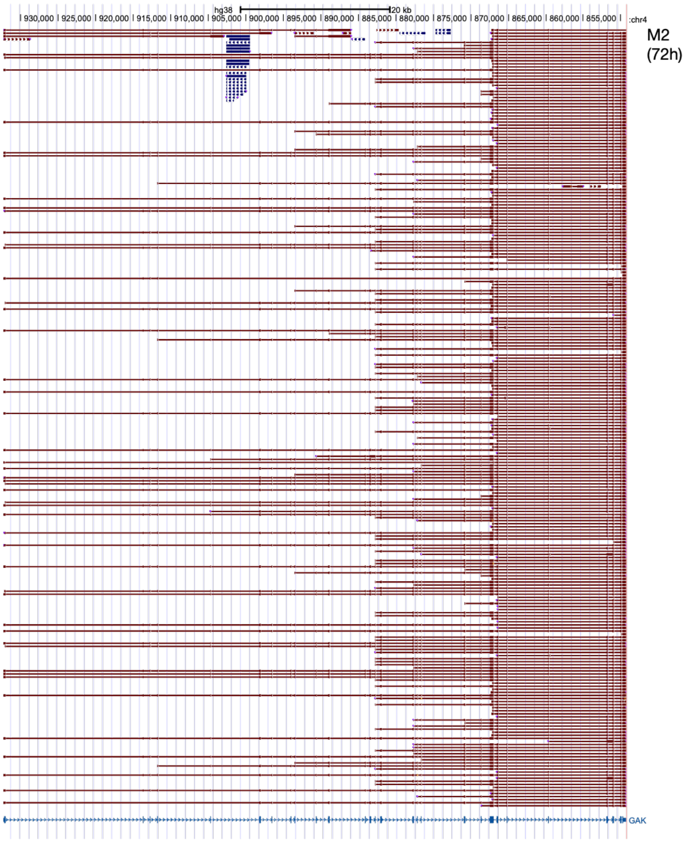
hg38
20 kb
930,000
925,000
920,000
915,000
910,000
905,000
900,000
895,000
890,000
885,000
880,000
875,000
870,000
865,000
860,000
855,000
:chr4
M2
(72h)
GAK

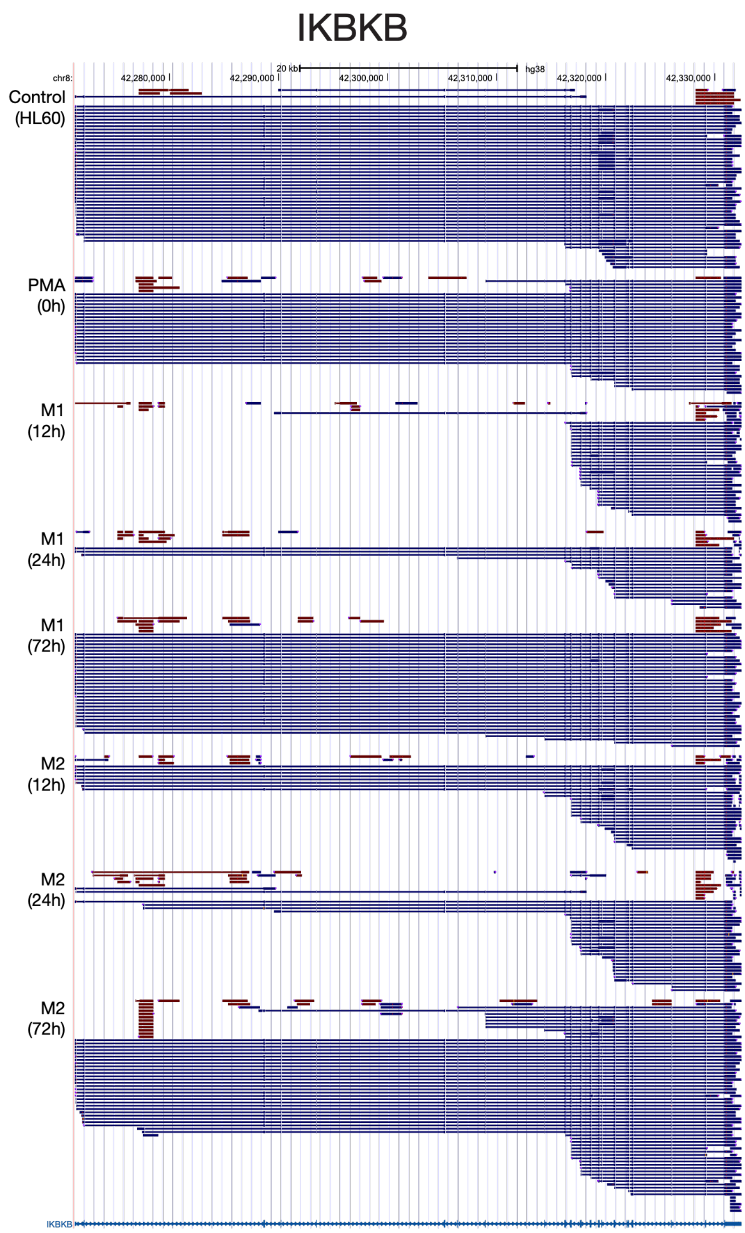

IKBKB
20 kb
hg38
chr8:
42,280,000
42,290,000
42,300,000
42,310,000
42,320,000
42,330,000
Control
(HL60)
PMA
(0h)
M1
(12h)
M1
(24h)
M1
(72h)
M2
(12h)
M2
(24h)
M2
(72h)
IKBKB


**B**

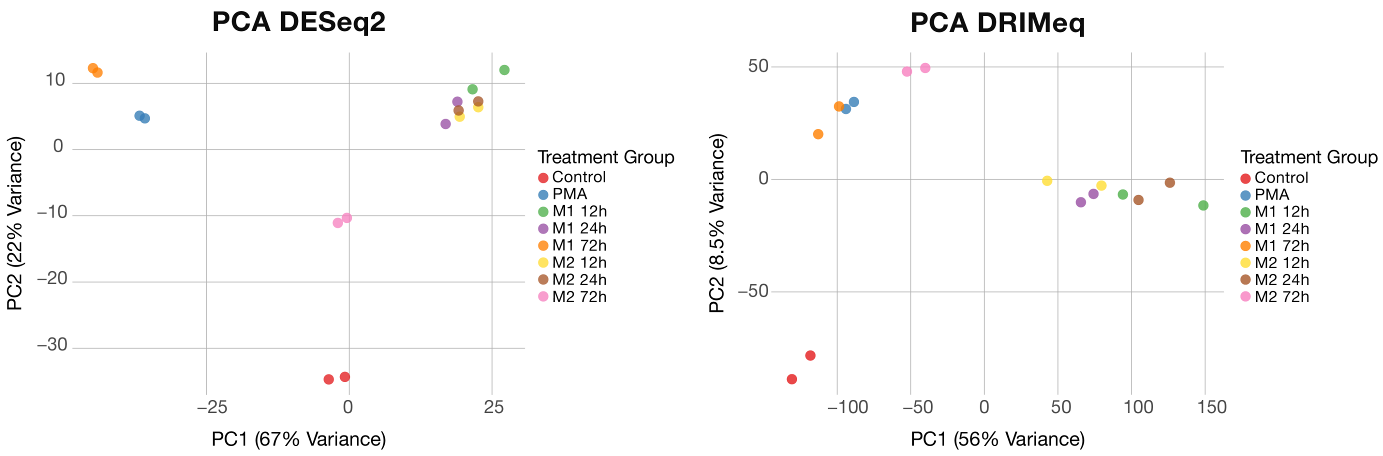

PCA DESeq2
PC2 (22% Variance)
10
0
−10
−20
−30
−25
0
25
PC1 (67% Variance)
Treatment Group
Control
PMA
M1 12h
M1 24h
M1 72h
M2 12h
M2 24h
M2 72h
PCA DRIMeq
PC2 (8.5% Variance)
50
0
−50
−100
−50
0
50
100
150
PC1 (56% Variance)
Treatment Group
Control
PMA
M1 12h
M1 24h
M1 72h
M2 12h
M2 24h
M2 72h

**Supplementary Figure 1.**
**(A)** Representative long-read RNA-seq alignments across the GAK and IKBKB loci in untreated HL-60 cells, PMA-treated cells, and M1- or M2-polarized cells collected at 12, 24, and 72 h. Individual reads illustrate condition- and time-dependent changes in transcript architecture, including the appearance and disappearance of truncated RNA species during macrophage polarization. **(B)** Principal-component analysis (PCA) of the same samples using DESeq2 gene-level abundance measurements (left) or DRIMSeq transcript-usage profiles (right). Samples segregate according to treatment and polarization state, demonstrating reproducible, condition-dependent changes in both gene abundance and transcript composition.

### Supplementary Figure 2

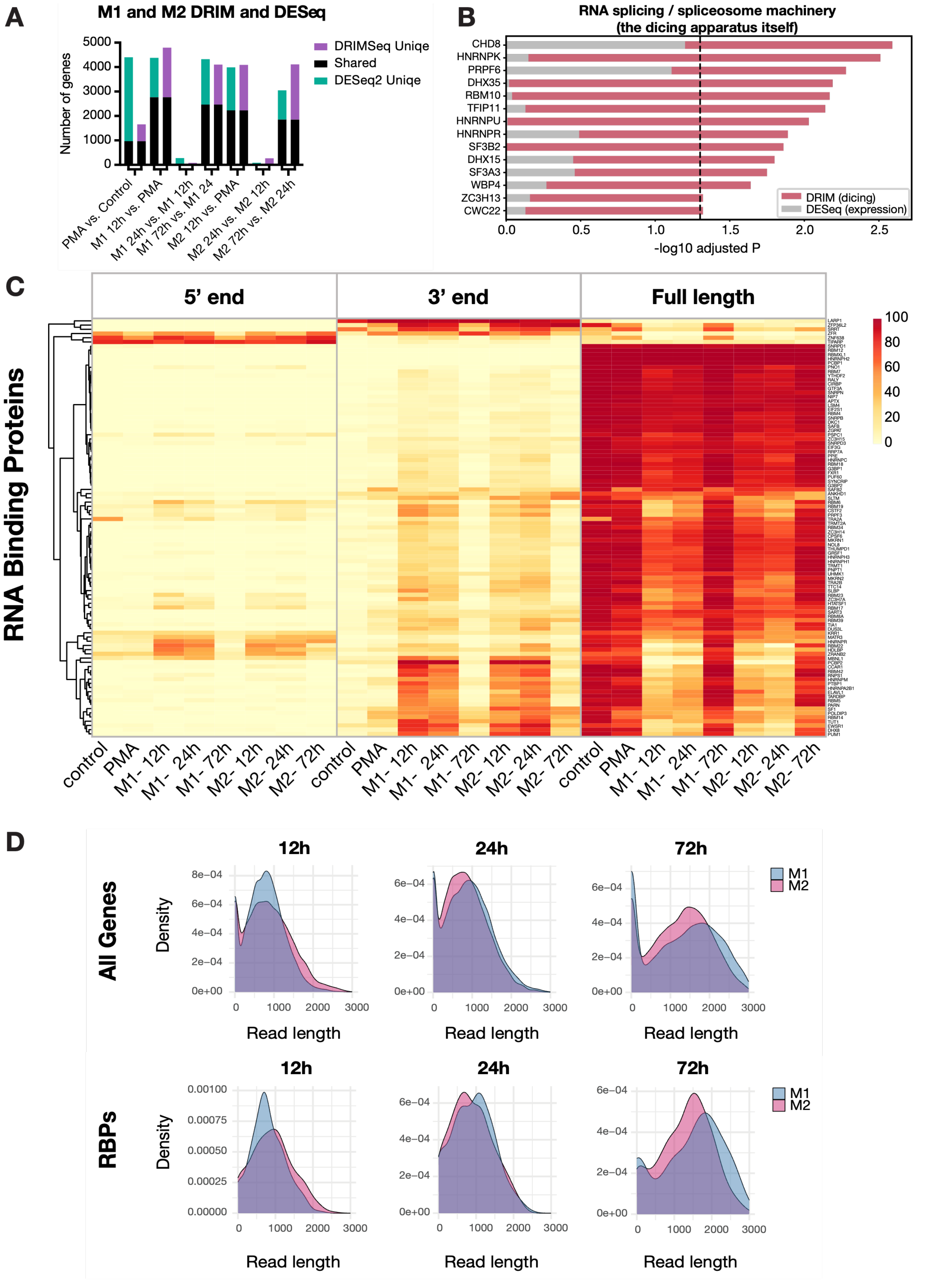

E

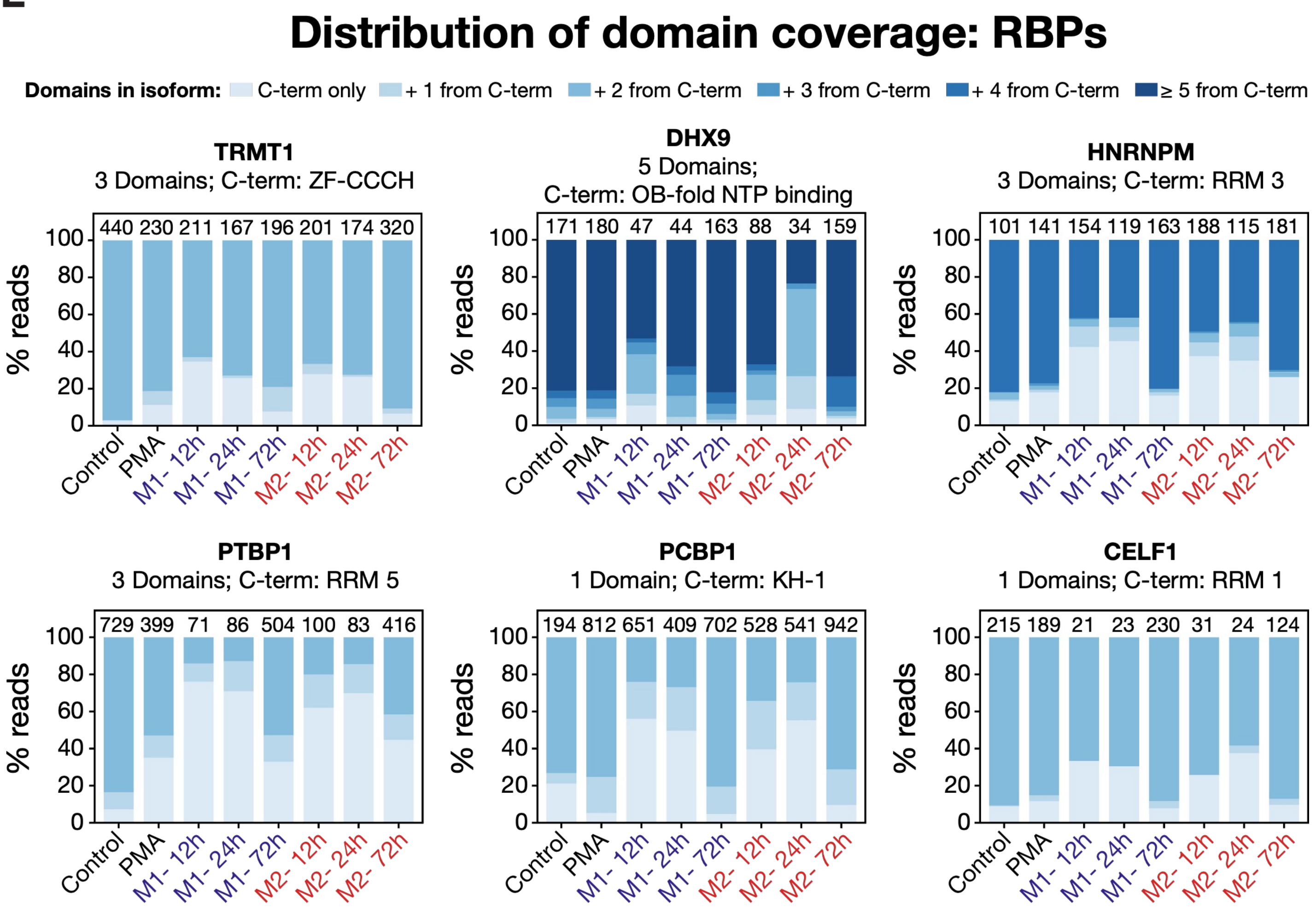

Distribution of domain coverage: RBPs
Domains in isoform: C-term only + 1 from C-term + 2 from C-term + 3 from C-term + 4 from C-term ≥ 5 from C-term
TRMT1
3 Domains; C-term: ZF-CCCH
440 230 211 167 196 201 174 320
DHX9
5 Domains;
C-term: OB-fold NTP binding
171 180 47 44 163 88 34 159
HNRNPM
3 Domains; C-term: RRM 3
101 141 154 119 163 188 115 181
PTBP1
3 Domains; C-term: RRM 5
729 399 71 86 504 100 83 416
PCBP1
1 Domain; C-term: KH-1
194 812 651 409 702 528 541 942
CELF1
1 Domains; C-term: RRM 1
215 189 21 23 230 31 24 124
% reads
0 20 40 60 80 100
Control PMA M1- 12h M1- 24h M1- 72h M2- 12h M2- 24h M2- 72h


F

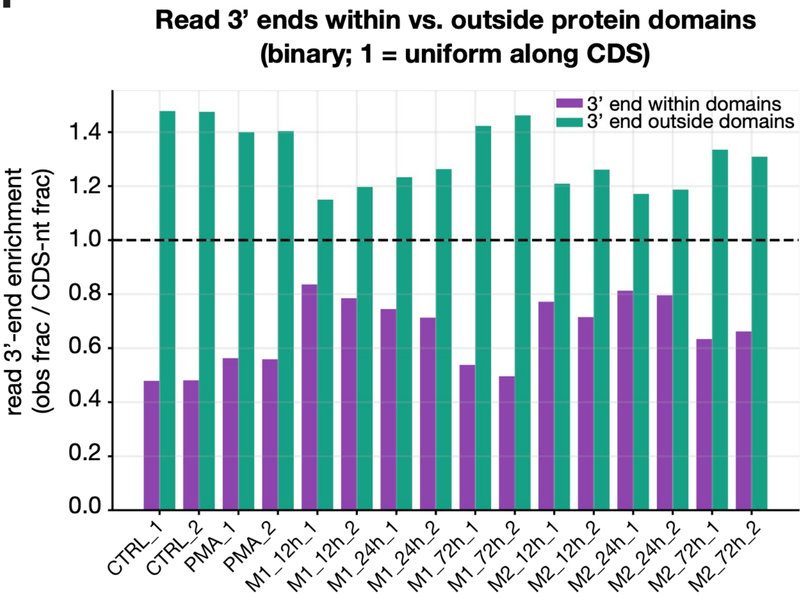

Read 3’ ends within vs. outside protein domains
(binary; 1 = uniform along CDS)
3’ end within domains
3’ end outside domains
read 3’-end enrichment
(obs frac / CDS-nt frac)
0.0 0.2 0.4 0.6 0.8 1.0 1.2 1.4
CTRL_1 CTRL_2 PMA_1 PMA_2 M1_12h_1 M1_12h_2 M1_24h_1 M1_24h_2 M1_72h_1 M1_72h_2 M2_12h_1 M2_12h_2 M2_24h_1 M2_24h_2 M2_72h_1 M2_72h_2

**Supplementary Figure 2.**
**(A)** Numbers of genes identified as significantly altered by DESeq2 and DRIMSeq across consecutive stages of differentiation, including PMA versus untreated control, PMA to 12 h polarization, and subsequent 12-to-24-h and 24-to-72-h transitions in the M1 and M2 lineages. Bars indicate genes uniquely detected by DESeq2 or DRIMSeq and genes shared between the two analyses. **(B)** Differential expression and dicing of genes associated with RNA splicing and spliceosome machinery. DRIMSeq and DESeq2 significance values are shown as –log10 adjusted *P* values, highlighting RNA-processing factors that undergo changes in transcript composition that are not equivalently reflected at the gene-expression level. **(C)** Heatmap showing the relative abundance of 5′, 3′, and full-length transcript classes for RNA-binding proteins across control, PMA, and M1/M2 polarization at 12, 24, and 72 h. RNA-binding proteins display dynamic and fate-dependent changes in transcript architecture similar to those observed for protein kinases. **(D)** Density distributions of 3′-isoform read lengths for all genes (upper panels) and RNA-binding proteins (RBPs; lower panels) in M1 and M2 cells at 12, 24, and 72 h, illustrating time- and fate-dependent differences in the extent of transcript truncation. **(E)** Domain coverage of representative RNA-binding proteins across differentiation states. Stacked bars indicate the fraction of reads retaining progressively larger numbers of protein domains counted from the C terminus; numbers above bars indicate the corresponding read counts. **(F)** Enrichment of read 3′ ends within versus outside annotated protein domains, normalized to the fraction of coding-sequence nucleotides belonging to each category (1 = expectation for a uniform distribution across the CDS). Transcript boundaries are preferentially enriched outside annotated domains, consistent with dicing occurring in inter-domain regions and preserving downstream domain architecture.

### Supplementary Figure 3

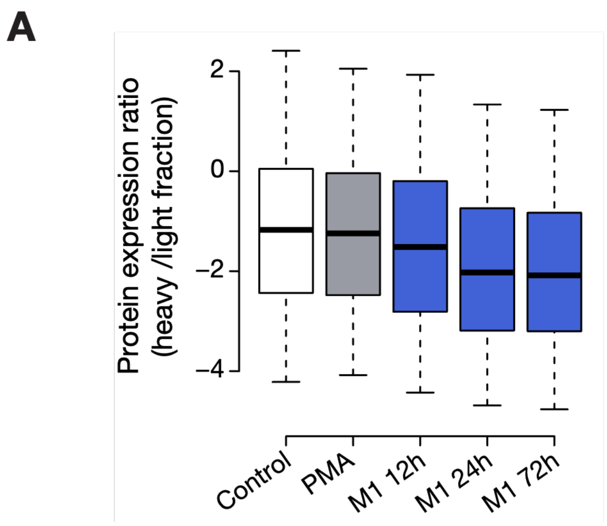


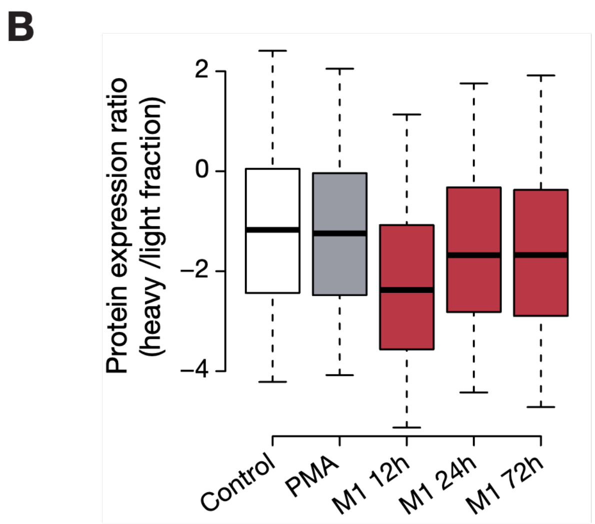


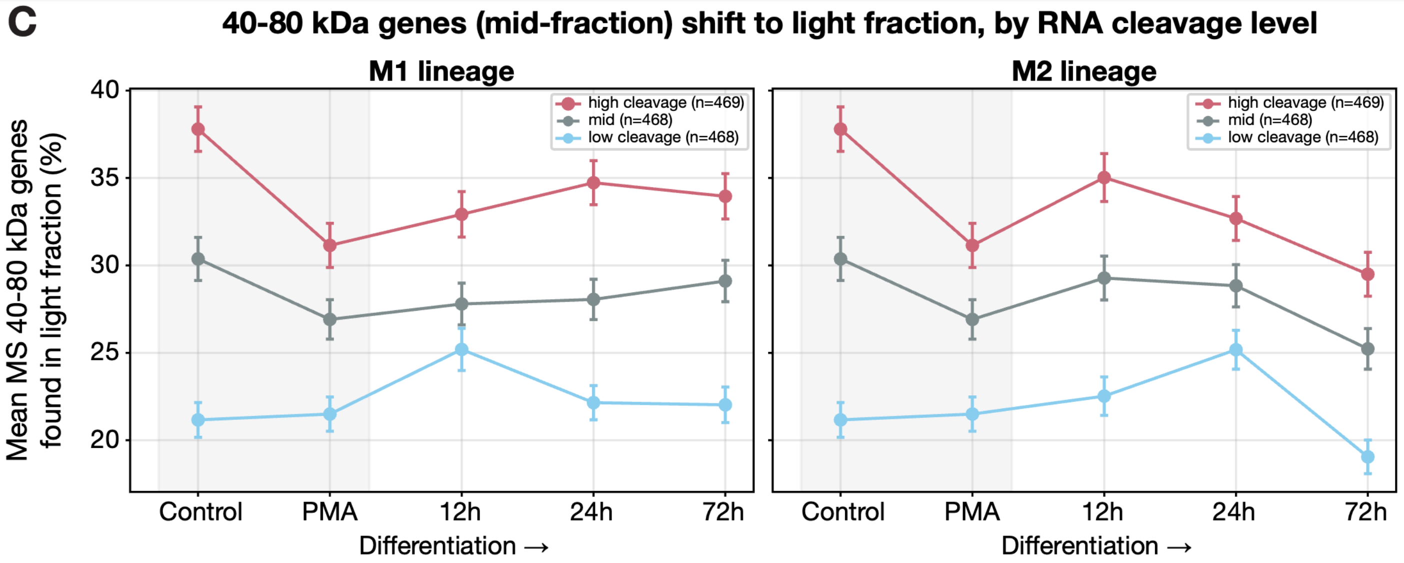

**D**

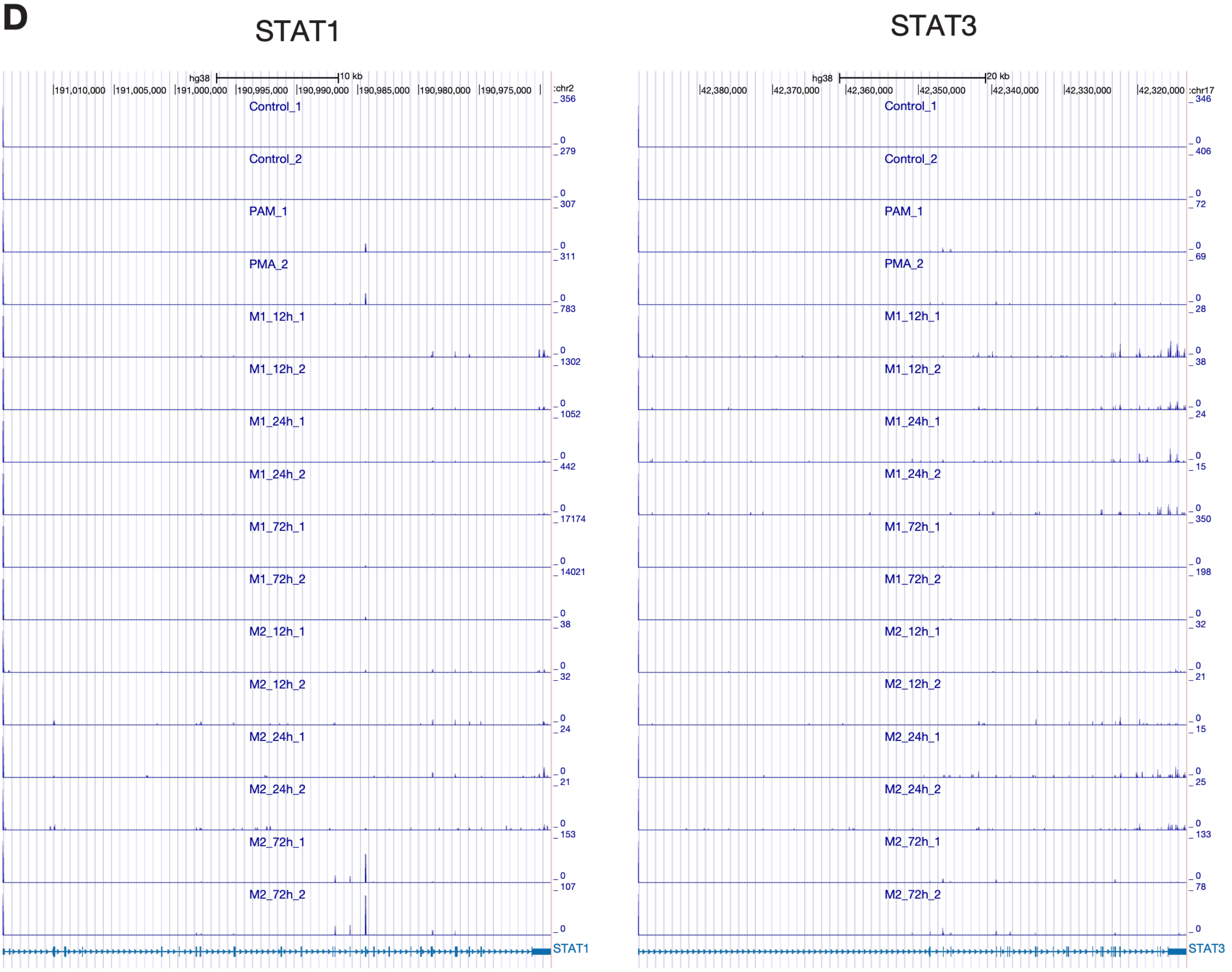
STAT1
STAT3
hg38
10 kb
20 kb
191,010,000
191,005,000
191,000,000
190,995,000
190,990,000
190,985,000
190,980,000
190,975,000
chr2
42,380,000
42,370,000
42,360,000
42,350,000
42,340,000
42,330,000
42,320,000
chr17
Control_1
Control_2
PAM_1
PMA_2
M1_12h_1
M1_12h_2
M1_24h_1
M1_24h_2
M1_72h_1
M1_72h_2
M2_12h_1
M2_12h_2
M2_24h_1
M2_24h_2
M2_72h_1
M2_72h_2
STAT1
STAT3

**E**

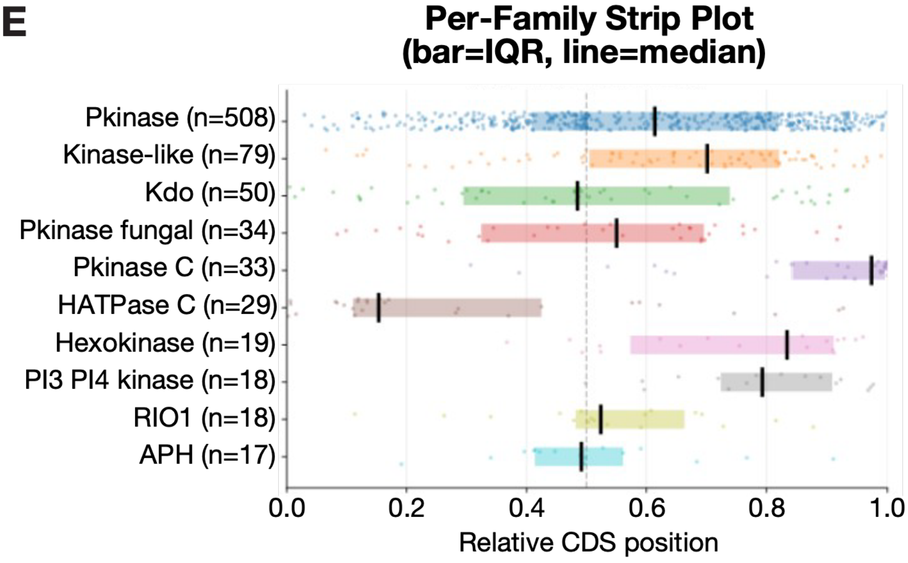
Per-Family Strip Plot
(bar=IQR, line=median)
Pkinase (n=508)
Kinase-like (n=79)
Kdo (n=50)
Pkinase fungal (n=34)
Pkinase C (n=33)
HATPase C (n=29)
Hexokinase (n=19)
PI3 PI4 kinase (n=18)
RIO1 (n=18)
APH (n=17)
0.0
0.2
0.4
0.6
0.8
1.0
Relative CDS position

**Supplementary Figure 3.**

**(A, B)** Distribution of heavy-to-light protein-expression ratios across control, PMA, and successive polarization time points in the M1 (**A**) and M2 (**B**) lineages. The temporal shift toward the light protein fraction follows the corresponding RNA-dicing dynamics, with distinct kinetics in M1- and M2-polarized cells. **(C)** Relationship between RNA-cleavage intensity and redistribution of proteins with canonical molecular weights of 40–80 kDa toward the light fraction. Genes were stratified into high-, intermediate-, and low-cleavage groups, and the mean percentage detected in the light fraction is shown across differentiation for the M1 and M2 lineages. Genes with greater RNA cleavage show a consistently greater shift toward lower-molecular-weight protein species. **(D)** 5’end of Long-read RNA-seq profiles of STAT1 and STAT3 across control, PMA, and M1/M2 polarization, showing dynamic and lineage-dependent remodeling of transcript architecture. **(E)** Relative coding-sequence position of annotated kinase-related protein domains across major domain families. Individual domains are shown according to their relative position along the CDS; bars indicate the interquartile range and vertical lines indicate the median. Many kinase-domain families are preferentially positioned toward the C-terminal portion of the coding sequence, providing an architecture in which RNA dicing can remove upstream regions while retaining downstream catalytic modules.

### Supplementary Figure 4

**A**

JAK1

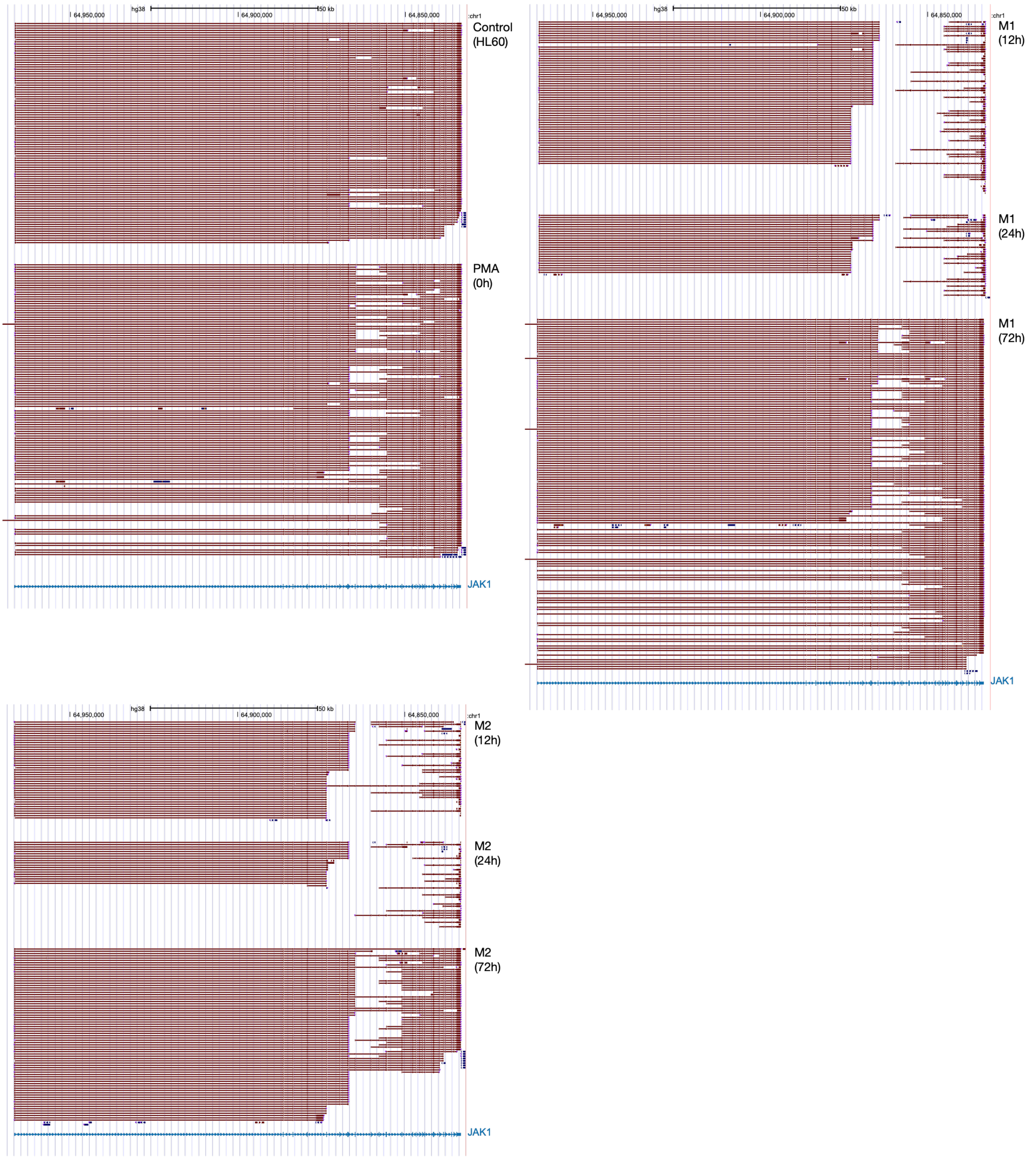

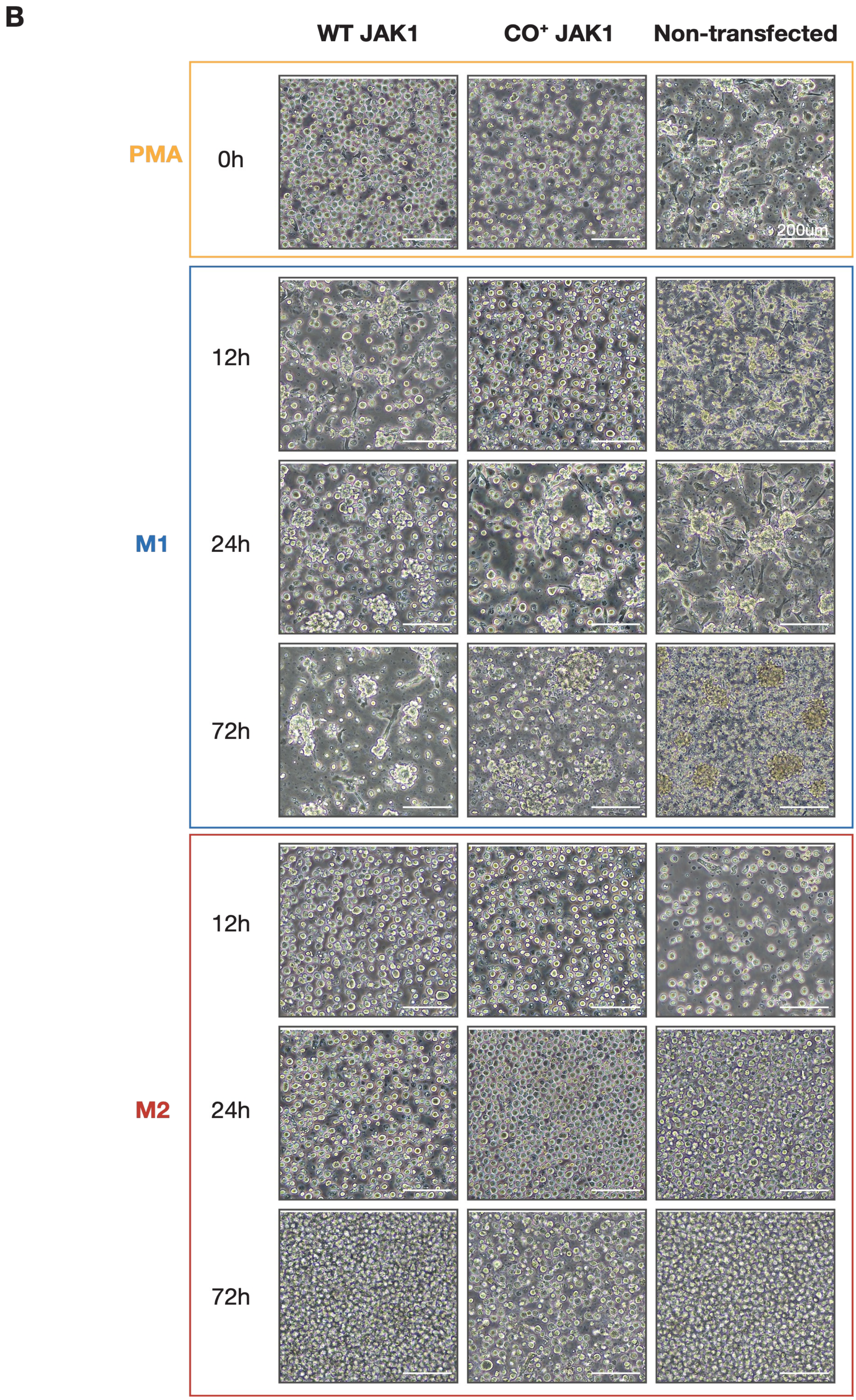
B
WT JAK1
CO⁺ JAK1
Non-transfected
PMA
0h
200um
M1
12h
24h
72h
M2
12h
24h
72h

C

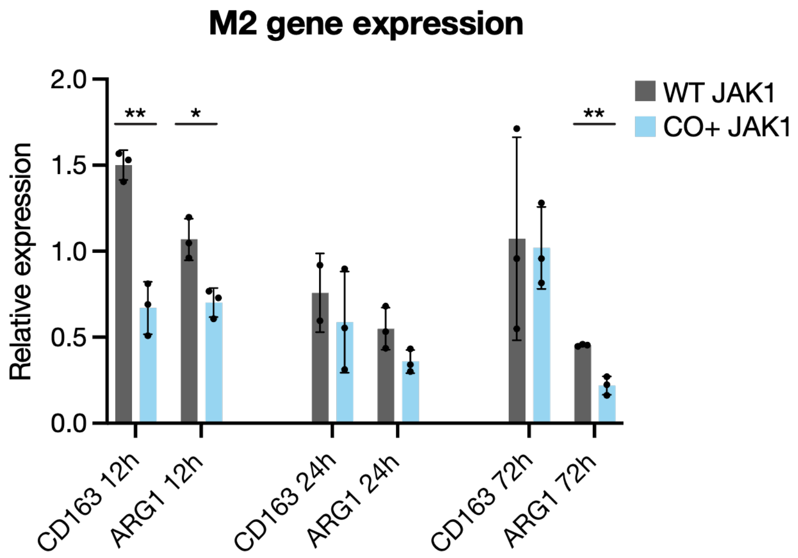

M2 gene expression
Relative expression
2.0
1.5
1.0
0.5
0.0
**
*
**
WT JAK1
CO+ JAK1
CD163 12h
ARG1 12h
CD163 24h
ARG1 24h
CD163 72h
ARG1 72h

**Supplementary Figure 4**

**(A)** Representative long-read RNA-seq alignments across the JAK1 locus in untreated HL-60 cells, PMA-treated cells, and M1- or M2-polarized cells at 12, 24, and 72 h. JAK1 displays pronounced transcript remodeling during early polarization followed by a return toward the PMA-associated transcript architecture at later time points. **(B)** Representative phase-contrast images of HL-60 cells expressing wild-type (WT) JAK1 or the synonymous codon-modified, dicing-impaired JAK1 allele (CO+), together with non-transfected cells, following PMA treatment and M1 or M2 polarization. WT JAK1 cells develop the characteristic clustered morphology associated with M1 differentiation, whereas this phenotype is reduced in cells expressing CO+ JAK1. Scale bar, 200 µm. **(C)** qRT-PCR analysis of the M2-associated markers CD163 and ARG1 in cells expressing WT or CO+ JAK1 at 12, 24, and 72 h following M2 induction. Disruption of JAK1 dicing produces comparatively modest effects on M2-associated gene expression relative to the pronounced effects observed during M1 polarization. $P < 0.05$ (*), $P < 0.01$ (**), two-tailed Student's *t*-test.

### Supplementary Figure 5

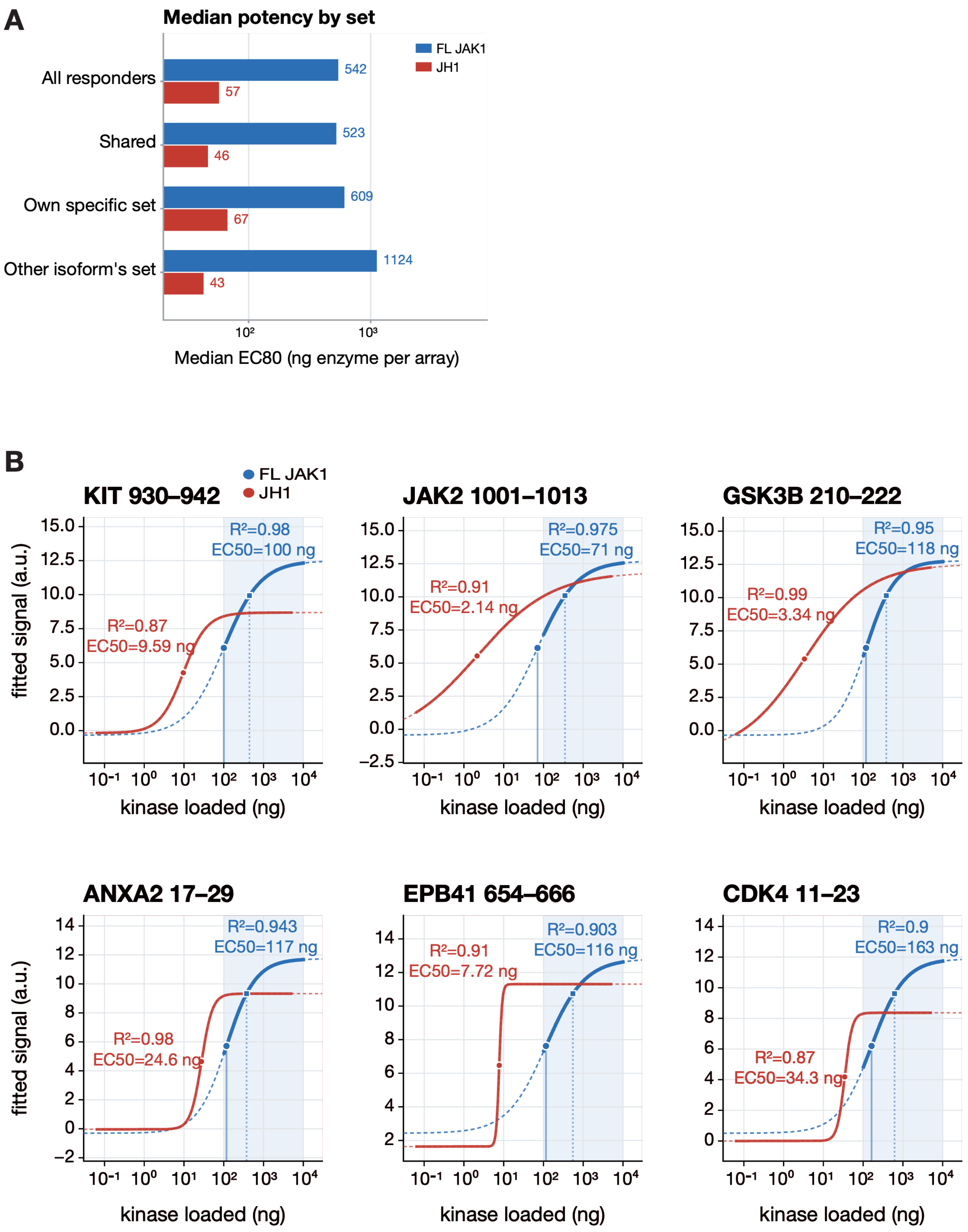

C

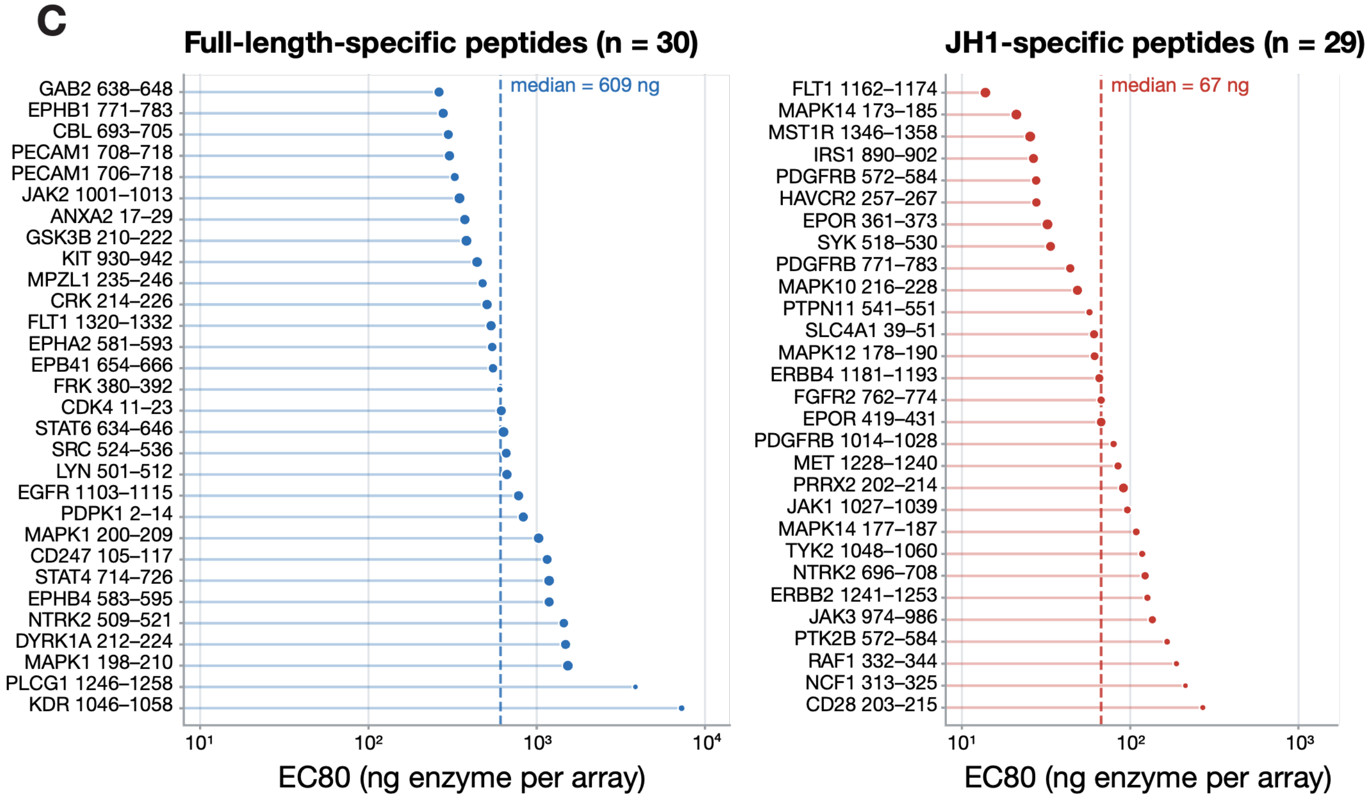
Full-length-specific peptides (n = 30)
median = 609 ng
GAB2 638–648
EPHB1 771–783
CBL 693–705
PECAM1 708–718
PECAM1 706–718
JAK2 1001–1013
ANXA2 17–29
GSK3B 210–222
KIT 930–942
MPZL1 235–246
CRK 214–226
FLT1 1320–1332
EPHA2 581–593
EPB41 654–666
FRK 380–392
CDK4 11–23
STAT6 634–646
SRC 524–536
LYN 501–512
EGFR 1103–1115
PDPK1 2–14
MAPK1 200–209
CD247 105–117
STAT4 714–726
EPHB4 583–595
NTRK2 509–521
DYRK1A 212–224
MAPK1 198–210
PLCG1 1246–1258
KDR 1046–1058
$10^1$
$10^2$
$10^3$
$10^4$
EC80 (ng enzyme per array)
JH1-specific peptides (n = 29)
median = 67 ng
FLT1 1162–1174
MAPK14 173–185
MST1R 1346–1358
IRS1 890–902
PDGFRB 572–584
HAVCR2 257–267
EPOR 361–373
SYK 518–530
PDGFRB 771–783
MAPK10 216–228
PTPN11 541–551
SLC4A1 39–51
MAPK12 178–190
ERBB4 1181–1193
FGFR2 762–774
EPOR 419–431
PDGFRB 1014–1028
MET 1228–1240
PRRX2 202–214
JAK1 1027–1039
MAPK14 177–187
TYK2 1048–1060
NTRK2 696–708
ERBB2 1241–1253
JAK3 974–986
PTK2B 572–584
RAF1 332–344
NCF1 313–325
CD28 203–215
$10^1$
$10^2$
$10^3$
EC80 (ng enzyme per array)

D

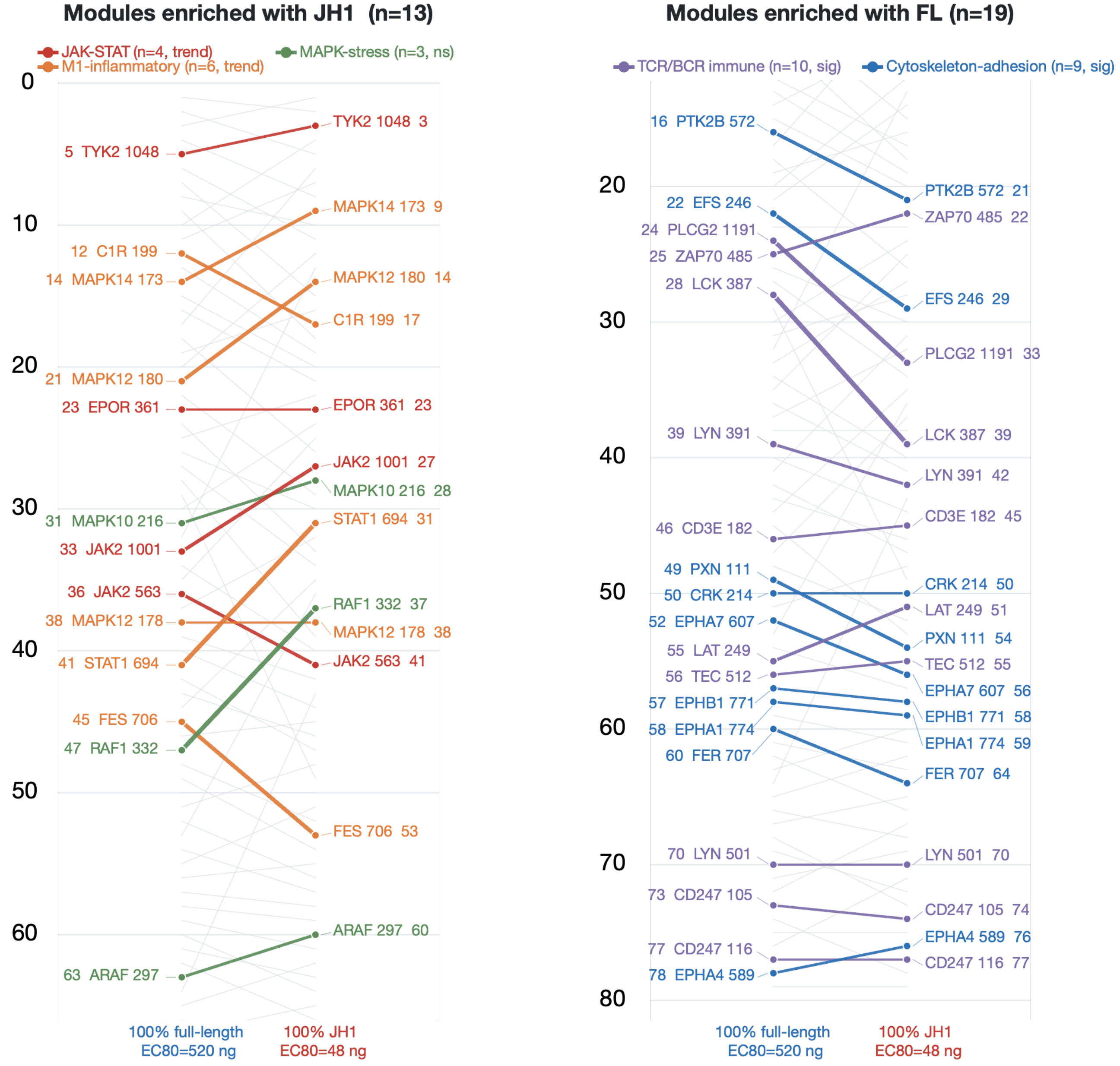
Modules enriched with JH1 (n=13)
JAK-STAT (n=4, trend)
M1-inflammatory (n=6, trend)
MAPK-stress (n=3, ns)
0
10
20
30
40
50
60
5 TYK2 1048
12 C1R 199
14 MAPK14 173
21 MAPK12 180
23 EPOR 361
31 MAPK10 216
33 JAK2 1001
36 JAK2 563
38 MAPK12 178
41 STAT1 694
45 FES 706
47 RAF1 332
63 ARAF 297
TYK2 1048 3
MAPK14 173 9
MAPK12 180 14
C1R 199 17
EPOR 361 23
JAK2 1001 27
MAPK10 216 28
STAT1 694 31
RAF1 332 37
MAPK12 178 38
JAK2 563 41
FES 706 53
ARAF 297 60
100% full-length
EC80=520 ng
100% JH1
EC80=48 ng
Modules enriched with FL (n=19)
TCR/BCR immune (n=10, sig)
Cytoskeleton-adhesion (n=9, sig)
20
30
40
50
60
70
80
16 PTK2B 572
22 EFS 246
24 PLCG2 1191
25 ZAP70 485
28 LCK 387
39 LYN 391
46 CD3E 182
49 PXN 111
50 CRK 214
52 EPHA7 607
55 LAT 249
56 TEC 512
57 EPHB1 771
58 EPHA1 774
60 FER 707
70 LYN 501
73 CD247 105
77 CD247 116
78 EPHA4 589
PTK2B 572 21
ZAP70 485 22
EFS 246 29
PLCG2 1191 33
LCK 387 39
LYN 391 42
CD3E 182 45
CRK 214 50
LAT 249 51
PXN 111 54
TEC 512 55
EPHA7 607 56
EPHB1 771 58
EPHA1 774 59
FER 707 64
LYN 501 70
CD247 105 74
EPHA4 589 76
CD247 116 77
100% full-length
EC80=520 ng
100% JH1
EC80=48 ng

E

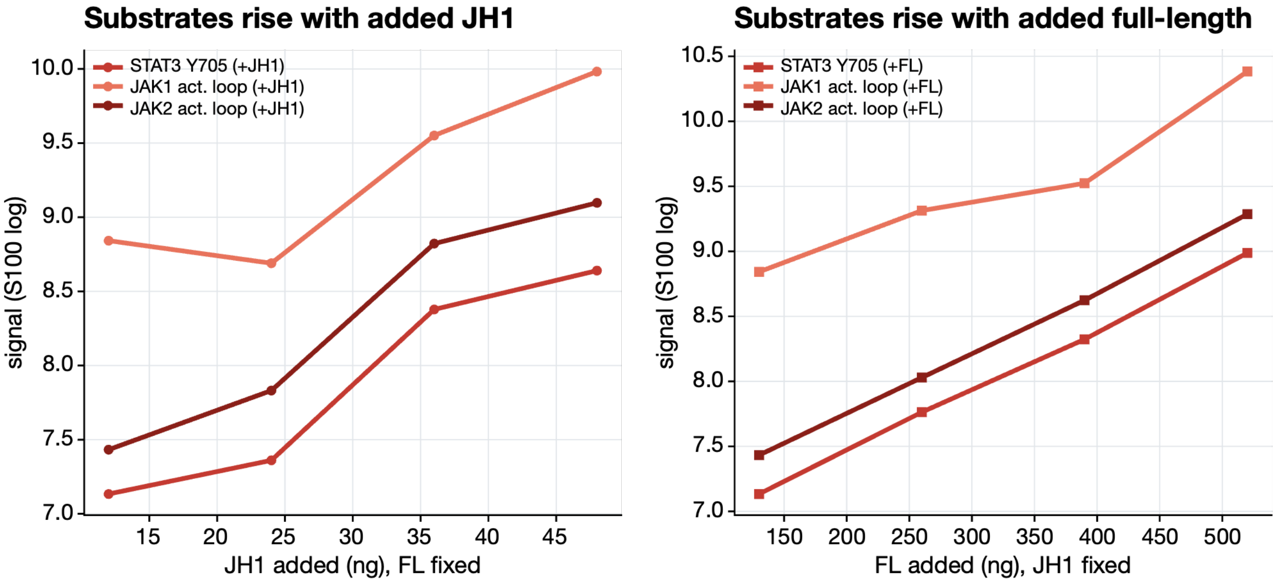

Substrates rise with added JH1
STAT3 Y705 (+JH1)
JAK1 act. loop (+JH1)
JAK2 act. loop (+JH1)
signal (S100 log)
10.0
9.5
9.0
8.5
8.0
7.5
7.0
15
20
25
30
35
40
45
JH1 added (ng), FL fixed
Substrates rise with added full-length
STAT3 Y705 (+FL)
JAK1 act. loop (+FL)
JAK2 act. loop (+FL)
signal (S100 log)
10.5
10.0
9.5
9.0
8.5
8.0
7.5
7.0
150
200
250
300
350
400
450
500
FL added (ng), JH1 fixed

**Supplementary Figure 5.**
**(A)** Median EC80 values for recombinant full-length JAK1 and the diced JH1 kinase module measured on the PamChip PTK peptide array. Values are shown for all responsive substrates, shared substrates, each isoform's preferential substrate set, and the substrate set preferentially targeted by the other isoform. Across all responsive peptides, diced JH1 reaches 80% of its maximal response at a substantially lower enzyme concentration than full-length JAK1. **(B)** Representative dose-response curves for selected peptide substrates phosphorylated by full-length JAK1 and diced JH1. Fitted responses, EC50 values, and goodness-of-fit ($R^2$) illustrate substrate-dependent differences in phosphorylation kinetics between the two kinase isoforms. **(C)** EC80 values for the 30 peptides preferentially phosphorylated by full-length JAK1 and the 29 peptides preferentially phosphorylated by diced JH1. Dashed lines indicate the median EC80 for each substrate group. **(D)** Rank-shift analysis of preferentially phosphorylated substrates between reactions containing 100% full-length JAK1 and 100% diced JH1. Substrates shifting toward diced JH1 are enriched for JAK–STAT, M1-inflammatory, and stress-MAPK-associated modules, whereas substrates preferentially retained by full-length JAK1 are enriched for immune-receptor and cytoskeletal/adhesion-associated modules. **(E)** Phosphorylation responses of selected STAT3, JAK1 activation-loop, and JAK2 activation-loop peptides during co-titration of the two JAK1 isoforms. Signals are shown as diced JH1 is increased at fixed full-length JAK1 (left) or full-length JAK1 is increased at fixed JH1 (right), illustrating that both isoforms remain catalytically active and contribute to substrate phosphorylation in mixed reactions.
